# Biostatistics and its impact on hazard characterization using in vitro developmental neurotoxicity assays

**DOI:** 10.1101/2022.10.18.512648

**Authors:** Hagen Eike Keßel, Stefan Masjosthusmann, Kristina Bartmann, Jonathan Blum, Arif Dönmez, Nils Förster, Jördis Klose, Axel Mosig, Melanie Pahl, Marcel Leist, Martin Scholze, Ellen Fritsche

## Abstract

In the field of hazard assessment, Benchmark concentrations (BMC) and their associated uncertainty are of particular interest for regulatory decision making. The BMC estimation consists of various statistical decisions to be made, which depend largely on factors such as experimental design and assay endpoint features. In current data practice, the experimenter is often responsible for the data analysis and therefore relies on statistical software without being aware about the software default settings and how they can impact the outputs of data analysis. To provide more insight into how statistical decision making can influence the outcomes of data analysis and interpretation, we have used case studies on a large dataset produced by a developmental neurotoxicity (DNT) in vitro battery (DNT IVB). Here we focused on the BMC and its confidence interval (CI) estimation, as well as on the final hazard classification. We identified five crucial statistical decisions experimenter have to face during data analysis: choice of replicate averaging, response data normalization, regression modelling, BMC and CI estimation, as well as choice of benchmark response levels. In addition, the strength of our data evaluation platform is the integration of endpoint-specific hazard classifications, including flagging systems for uncertain cases, which none of the so far existing statistical data analysis platforms provide. The insights gained in this study demonstrate how important fit-for-purpose, internationally harmonized and accepted data evaluation and analysis procedures are for an objective hazard classification.

## 1 Introduction

In 2007, the National Research Council (NRC) of the United States proposed a new strategy for toxicity testing in the 21^st^ century centering around a shift from *in vivo* experiments in animals to mechanism-based *in vitro* testing (NRC, 2007). Since then, major advances in the field of in vitro toxicology have been made, including development and establishment of medium and high throughput screening (HTS) assays, as well as bioinformatics tools for data generation, management and analysis (Leist et al., 2014; Wheeler et al., 2015; Villeneuve et al., 2019). These efforts are contributing to next generation risk assessment (NGRA), which aims at using new approach methods (NAMs) for exposure-based, hypothesis-driven risk assessment without the generation of new animal data (Li et al. 2021; Dent et al. 2021; Palloca et al. 2022).

Typically, an in vitro HTS test system produces hazard data for a relatively large number of test concentrations and thus makes it most suitable for concentration-response regression modelling. This statistical approach allows the interpolative estimation of a concentration value at a given effect level (effect or inhibitory concentration), and of particular regulatory interest is hereby the benchmark concentration (BMC) and its associated uncertainty, expressed as lower limit of a one-sided 95% confidence interval (BLL). A BMC is considered as lowest concentration of the test compound that produces a pre-defined small “relevant” change to the control reference’s response level, and a consequence, the benchmark response (BMR) value should be as “close as possible” to the control response.

In vitro test systems represent a huge variety of different types of assays, from cell-free, cell and tissue-based methods up to multi-response organoid systems, and as consequence, concentration-response data between these systems vary enormously with respect to their test-specific experimental designs, data variability, dynamic ranges and concentration-response pattern. Unique to HTS systems is also that assay outputs are produced in microplate multi-well readers, with concentration-response data from the same concentration and experiment are considered to reflect technical (intra-replicate) variation and data from repeated experiments more indicative for “biological” (between-study) variation. These hierarchical data are usually simplified by using an average response value per test concentration and experiment (replicate average) as statistical unit for the concentration-response analysis, with the argument that the BMC and BLL estimation should reflect mainly biological and between-study variability.

The BMC estimation consists of various statistical decisions to be made in the concentration-response analysis, which dependent largely on the experimental design, the concentration-response data and assay endpoint features, and which require statistical knowledge that is usually only warranted by experienced biostatisticians. In current data practice, the experimenter is often responsible for the data analysis and therefore relies on statistical software without being aware about the software default settings and how they can impact the outputs of data analysis (Jensen *et al*., 2020). Existing guidelines for concentration response data analysis are often too general (OECD, 2006; EFSA, 2016), and no clear consensus on a common and standardized biostatistical method for *in vitro* toxicity data have been achieved (Wheeler et al., 2015; Sand et al., 2017).

To provide more insight into how statistical decision making can influence the outcomes of data analysis and interpretation, we have used case studies on a large dataset produced by a developmental neurotoxicity (DNT) in vitro battery (DNT IVB; Masjosthusmann et al. 2020, Crofton and Mundy 2021). In this DNT IVB, 148 compounds were tested across up to ten test methods representing the neurodevelopmental key events (KE) of neural progenitor cell (NPC) proliferation, migration of neural crest and radial glia cells, neurons and oligodendrocytes, neuronal differentiation, neurite outgrowth of peripheral and central nervous system neurons, as well as oligodendrocyte differentiation, and accomplished by various endpoints measuring cell viability and cytotoxicity (Masjosthusmann et al. 2020). Some of the DNT-specific endpoints are derived from primary and organotypic cultures, and thus more prone to a data variability typically observed in animal studies. Here we focused on the BMC and its confidence interval (CI) estimation, as well as the final hazard classification. For this purpose, we identified five crucial statistical decisions the experimenter have to face during the data analysis (Figure 1):

**Figure 1:**
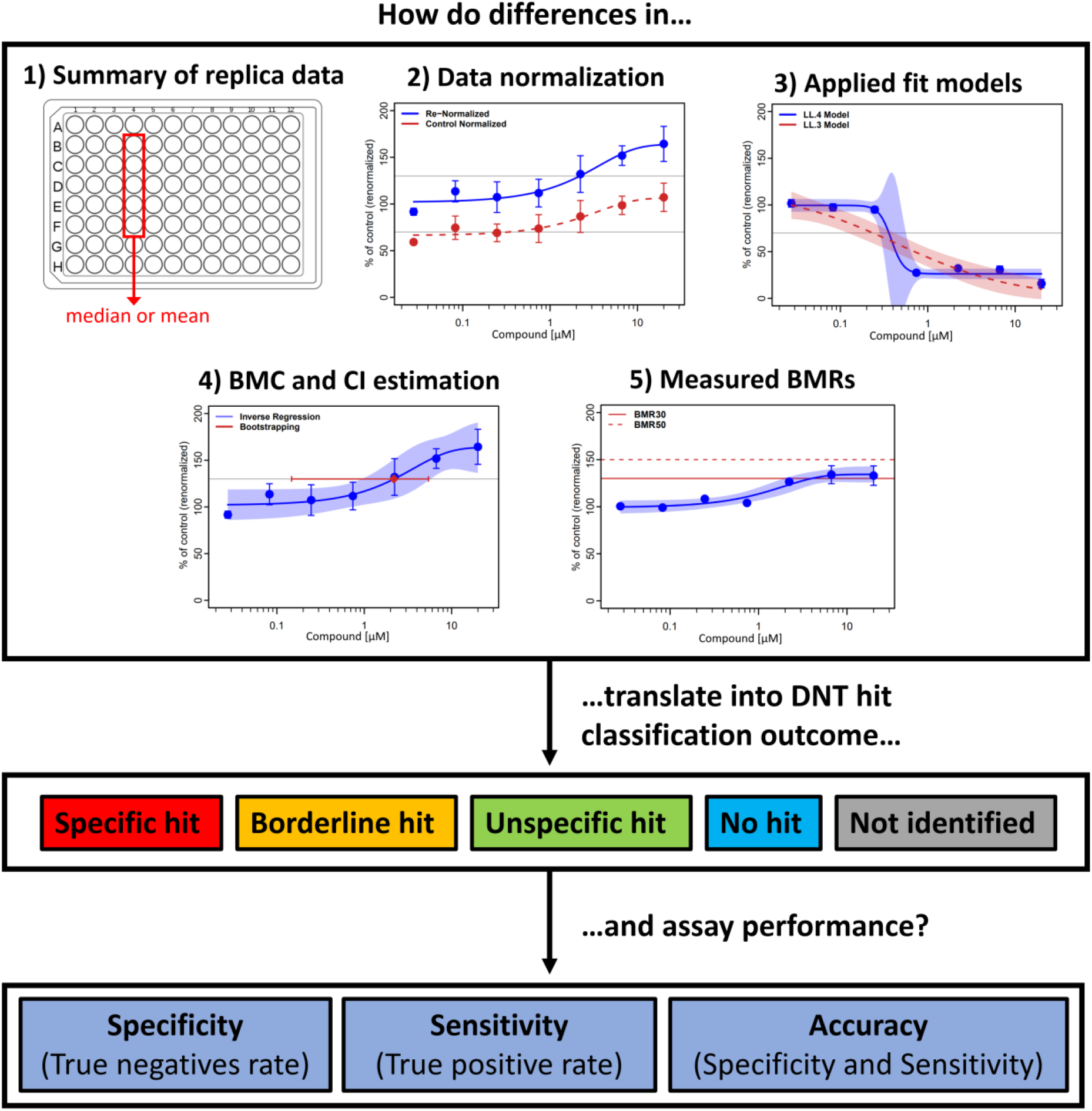
Study Overview. Several biostatistical data analysis and evaluation steps were analysed for their impact on a BMC estimation and subsequent hazard characterization from developmental neurotoxicity (DNT) data: i) how to average replicate responses from an experiment, ii) how to normalize concentration-response data, iii) how to describe concentration-response data by regression modelling, iv) how to estimate a benchmark concentration (BMC) and its uncertainty, and v) which benchmark response (BMR) level to select. Changes between statistical methods were recorded for 148 compounds tested on up to 22 assay endpoints, and their impact translated into the compound’s DNT hit classification and the predictivity performance of the overall assay battery.

i. Statistical unit: shall the median of all replicate responses of an experiment be used, which makes no assumption to the data and thus reduces the negative impact of potential data outlier, or the replicate mean, which has a higher certainty but assumes a symmetric distribution of the replicate responses, and if violated, can lead to a biased estimation of the replicate mean?
ii. Response data normalization: shall the responses of an experiment always be normalized to the control’s response even if the exposure responses provide clear evidence against the use of control data, or shall in that case the “control reference” be estimated directly from the responses of the exposures (“re-normalization”, Krebs et al., 2018)?
iii. Regression model: shall the concentration-response data always be described by the same and supposedly flexible mathematical model, or is it better to use several models and either subsequently select the best model by means of goodness-of-fit criteria (“best-fit method”, Scholze *et al*., 2001) or estimate an average of all model fits (“model averaging”, Claeskens et a, 2008)?
iv. Uncertainty of a BMC estimation: shall the confidence level of a BMC be calculated by a simple and commonly used statistical approximation technique (“Delta method”, Cox, 1990) which is known to be inaccurate (Moerbeek, Piersma and Slob, 2004), or by alternative approaches such as bootstrapping or inverse regression (Jensen, Kluxen and Ritz, 2019)?
v. Benchmark response (BMR): shall a response level most close to the control reference be selected, which might not always be applicable for the statistical concentration-response analysis and thus might fail to provide a reliable BMC estimation, or a higher BMR, which guarantees a statistically more robust BMC estimation but might fail for compounds that has produced weak responses below the intended BMR?

We designed a standard data evaluation protocol (“standard protocol”) which we used as reference to alternative statistical methods, so that their BMCs and confidence intervals (CIs) estimated to the same DNT IVB data could be compared. The statistical methods to be changed were chosen along the questions outlined in (i) to (iv). This was supplemented by measuring their impact on hazard alerts derived from hit classifications, which separate cytotoxic concentration ranges from the respective BMC of the specific DNT endpoint, and by measuring their impact on the DNT IVB’s capability on predicting DNT adversity in terms of specify, sensitivity and accuracy.

## 2 Methods

### 2.1 DNT data

All concentration response data used in this study are from a DNT *in vitro* battery of 8 assays with 22 endpoints, in which a total of 148 compounds were tested. 120 compounds were tested across all assays, while 28 compounds were tested in at least 2 assays. Fourteen assay endpoints represent major key neurodevelopmental processes, and 8 endpoints measure general cell viability and cytotoxicity (Table 1). This DNT *in vitro* battery was developed in collaboration with EFSA with the aim to advance the application of *in vitro* DNT testing for regulatory purposes. The term “BMC” was used equally for data from DNT-specific, cytotoxicity and viability endpoints.

**Table 1:** Test Assays. An overview over the assays and their key characteristics, including the cell model, assays endpoint and chemical exposure time (in brackets), the BMR that was used for the classification, the number of independent experiments as well as the replicates per experiment. Unspecific endpoints (cell toxicity and viability) are highlighted in cursive. Cytotoxicity and cell viability assay endpoints were used as reference for NPC2, NPC3 and NPC5.

| Assay | Cell model | Endpoint [exposure time] | BMR [%] | Independent experiments | #Replicate experiment | # of Compounds<sup>2</sup> |
| --- | --- | --- | --- | --- | --- | --- |
| NPC1 | neural progenitor cells | proliferation NPC1 [72h] | 30 | 3 to 5 | 5 | 123 |
|  |  | proliferation by area [72h] | 30 |  |  | 117 |
|  |  | *viability NPC1 [72h]* | 30 |  |  | 123 |
|  |  | *cytotoxicity NPC1 [72h]* | 10 |  |  | 115 |
| NPC2 | neural progenitor cells | migration distance radial glia NPC2a [72h] | 10 | 3 to 5 | 5 | 123 |
|  |  | migration distance radial glia NPC2a [120h] | 10 |  |  |  |
|  |  | migration distance neurons NPC2b [120h] | 30 |  |  |  |
|  |  | migration distance oligodendrocytes NPC2c [120h] | 30 |  |  |  |
| NPC3 | neural progenitor cells | neuronal differentiation NPC3 [120h] | 30 | 3 to 5 | 5 | 123 |
|  |  | neurite length NPC4 [120h] | 30 |  |  |  |
|  |  | neurite area NPC4 [120h] | 30 |  |  |  |
| NPC5 | neural progenitor cells | oligodendrocyte differentiation NPC5 [120h] | 30 | 3 to 5 | 5 | 123 |
| NPC2-5<sup>1</sup> | neural progenitor cells | *cytotoxicity NPC2-5 [72h]* | 10 | 3 to 5 | 5 | 123 |
|  |  | *cell number [120h]* | 30 |  |  | 123 |
|  |  | *cytotoxicity NPC2-5 [120h]* | 10 |  |  | 122 |
|  |  | *viability NPC2-5 [120h]* | 30 |  |  | 123 |
| UKN2 | hiPSC-derived neural crest cells | Migration UKN2 [24h] | 25 | 3 | 6 | 70<sup>3</sup> |
|  |  | *Viability UKN2 [24h]* | 10 |  |  |  |
| UKN4 | Luhmes cells | Neurite Area UKN4 [24h] | 25 | 3 | 6 | 75<sup>3</sup> |
|  |  | *Viability UKN4 [24h]* | 25 |  |  |  |
| UKN5 | hiPSC-derived sensory neurons | Neurite Area UKN5 [24h] | 25 | 3 | 6 | 71<sup>3</sup> |
|  |  | *Viability UKN5 [24h]* | 25 |  |  |  |

<sup>1</sup>Cytotoxicity and cell viability assay endpoints were used as classification reference for NPC2, NPC3 and NPC5.
<sup>2</sup>Shown here is the number of compounds that have concentration response information with at least 5 concentrations
<sup>3</sup>A total of 140 compounds were tested in these assays. However, some compounds were only tested in a high concentration. If the high concentration was negative in the assay, no concentrations response testing was performed.

Depending on the assay, fluorescent readouts using a multiplate reader or fluorescence and brightfield imaging with subsequent artificial intelligence-based image analysis (Schmuck et al., 2015; Förster et al., 2021) was performed as endpoint assessment. Each compound was tested in at least three independent experiments and eight concentrations per experiment, with 5-6 controls and 5-6 replicates per experiment. An overview of the assays, the cell model and the respective endpoints is given in Table 1, and more detailed information about the assay-specific experimental testing procedures and test outcomes is provided in Masjosthusmann *et al*. (2020).

The BMR for each endpoint was derived from the between experimental variability as the coefficient of variation of median plate medians (after normalization) measured at the lowest test concentration and across all independent experiments (Masjosthusmann *et al*., 2020). To achieve a better comparability across the endpoints, the BMRs were then rounded to the next higher value, resulting into three BMRs: a 10% change was selected for endpoints from the NPC2a and NPC1-5 cytotoxicity assay (BMR10), and a 30% change for endpoints from the NPC1, NPC2a, NPC2b, NPC3-5 and NPC1-5 viability assay (BMR30). For all UKN assays a 25% change was decided (BMR25), and for the viability of the UKN2 a BMR10 was chosen. NPC Assays were conducted with three to five independent experiments and 5 replicates each, UKN Assays with three independent experiments and 6 replicates each (Table 1).

### 2.2 Data Evaluation Platform

For data processing and evaluation, the R package *drc* (R Core Team 2019, Ritz et al. 2019) was extended and optimized for the use of data from multi-well plate experiments. The biostatistical data evaluation software is freely available as open source under the name CRStats (github.com/ArifDoenmez/CRStats), an interactive R Markdown document is available and can freely be assessed for use. All for the comparative study relevant mayor modules are displayed as workflow diagram in Figure 2, starting from minimal data requirements for a BMC estimation (2.2.1) up to the endpoint-specific hazard classification module (2.2.8). The individual modules are explained in more detail below, with the module number in Figure 2 referring to the number of the subsection. From module 2.2.3 onwards we defined a standard protocol for the evaluation of DNT IVB data, with the following statistical methods chosen: (i) average replicate per experiment estimated by median (2.2.3), (ii) control-normalization followed by re-normalization (2.2.4), (iii) application of several mathematical models to find the ‘best fit’ regression model for a BMC estimation (2.2.6), (iv) CI estimation of the BMC by inverse regression (2.2.7), and (v) selecting endpoint specific BMRs for the hazard classification as outlined in Table 1. This standard setup is shown on the left side of the modules (blue), and all alternative methods that we considered in this study are listed on the right side under “changes” (orange).

**Figure 2:**
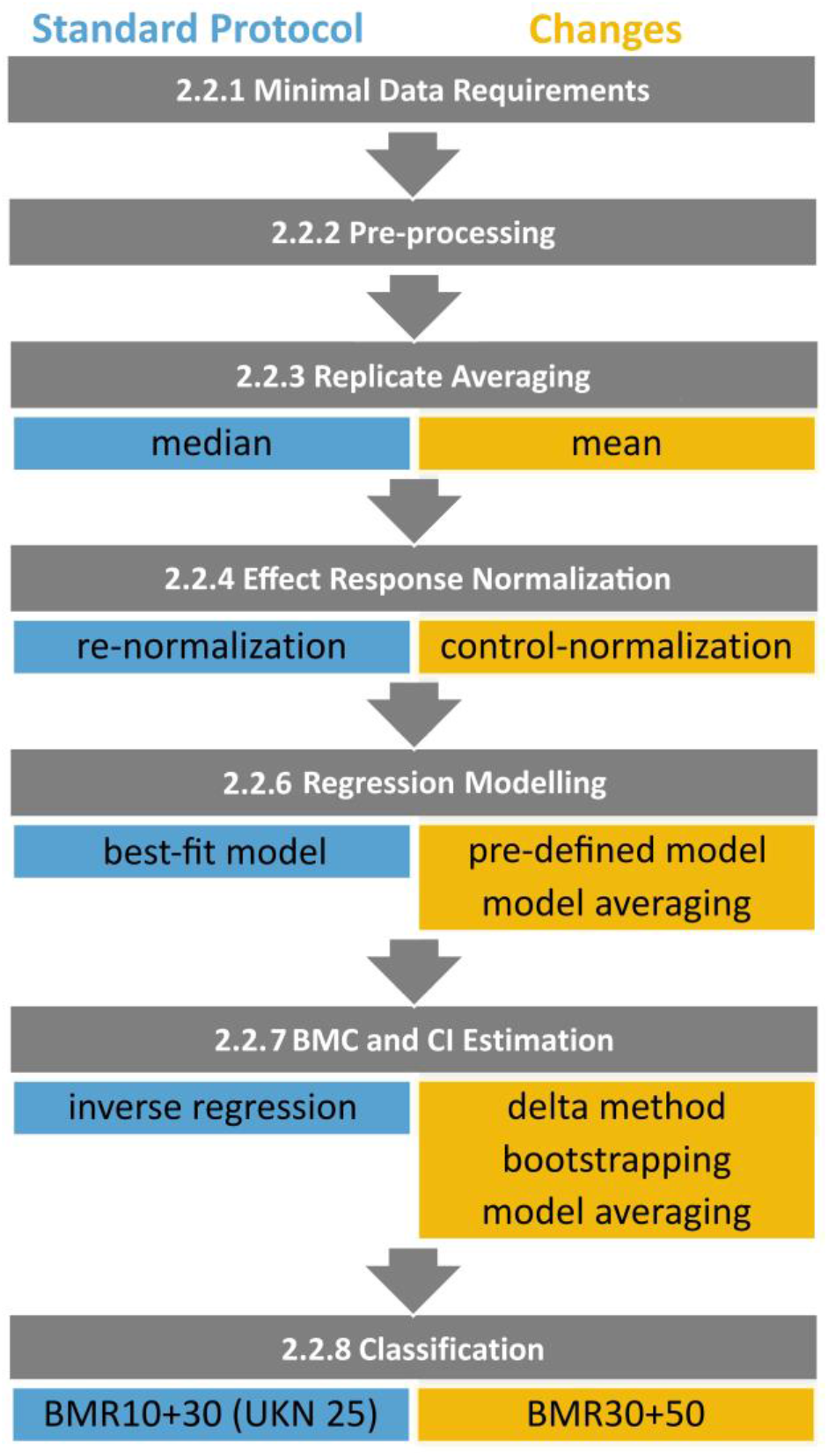
R Evaluation Pipeline Workflow. The R workflow is depicted with subsequent data evaluation steps from top to bottom. Grey boxes indicated the mayor data processing and evaluation steps with reference to the material and methods section in which they are described. Mayor key methods are depicted as coloured boxes below the according step. The blue methods (left) are the ones used for the standard protocol, while changes of one of the standard protocol methods (depicted in orange, right) are used to create the alternative protocols (i.e. for one alternative protocol the statistical unit for replicate averaging was changed from median to mean, while all other key methods remained the same as in the standard protocol, so that only the impact of this particular change can be monitored).

#### 2.2.1 Minimal data requirements

Data were accepted for data analysis only if the following three minimal data requirements were fulfilled: (i) at least two replicas per concentration are available, otherwise all readouts from this concentration were excluded, (ii) at least five concentrations per experiment have provided readouts otherwise the whole experiment was excluded, (iii) at least two control readouts are available otherwise the whole experiment was excluded.

#### 2.2.2 Pre-processing

CRSTATS uses different assay-specific pre-processing steps in order to obtain a single response value for each well. For example, the neuronal differentiation in the NPC3 assay is calculated as the number of neurons divided by the total number of cells with a nucleus:

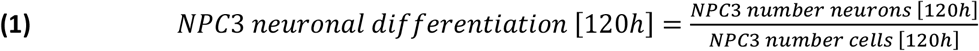

All assay specific pre-processing methods that are currently implemented in CRSTATS are listed in Table S1.

#### 2.2.3 Replicate averaging

The average assay response for controls and treatments from the same experiment was either estimated by the arithmetic mean or by the median. The variability between replicates was calculated as standard deviation (SD; for the mean replicate) or as median absolute deviation (MAD; for the median replicate). Outlier detection procedures were not applied and data points from wells where technical problems were known or obvious were excluded from the data analysis.

#### 2.2.4 Effect data normalization

CRSTATS offers different normalization methods which allows the translation of pre-processed effect data into relative values. For this study, we used the following two methods:

i. Control normalization: effect responses are normalized to the mean or median of the solvent controls as

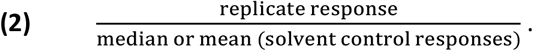
ii. Control re-normalization: normalized effect responses (Equation 2) are further normalized by a mean value that has been estimated by regression modelling at the lowest test concentration, i.e.

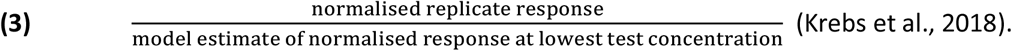

#### 2.2.5 Significance analysis

The presence of at least one exposure concentration that had produced an effect response which differs statistically significantly from the responses of all remaining exposures is a crucial factor in the hazard classification method (2.2.8). To account for that, significant differences between treatment means were identified by using the Tukey Honest Significant Differences test (alpha=5%, two-sided) (Tukey HSD; Yandell, 1997), with hypothesis testing conducted on normalized replicate averages from at least three independent experiments. As an average control value was always set to 100% (2.2.4), controls were excluded from the significance analysis. Data provided no evidence against the Gaussian assumption.

#### 2.2.6 Concentration-response regression analysis

The R packages *drc* (Ritz *et al*., 2015) and *bmd* (Jensen *et al*., 2020) were used for regression analysis and the estimation of a BMC and its associated uncertainty. The *drm* function fits a pre-defined regression model to the concentration-response data, with several options implemented to provide more flexibility for the estimation method. A large number of mathematical nonlinear regression functions was applied to the same data set (Table S2), and the best fitting model then selected on basis of the Akaike’s Information Criterion (AIC) (“best fit method”, Scholze et al., 2009; Portet, 2020). AIC is commonly used to compare the relative goodness-of-fit among different models and to then choose the model of best predictive power by balancing data support against model complexity. As all effect endpoints in this study are continuous, the estimation method of ordinary least-squares (OLS) was used. OLS relies on two assumptions, i.e. (i) effect data (here replicate average) follow a symmetrical distribution, and (ii) variance homogeneity across all treatment groups. Both assumptions were checked prior to data analysis on basis of pooled endpoint-specific data from all experiments: data variability differed in average by maximally 20% between the treatment groups, with the highest variability often occurring at highest test concentration, and no overall clear evidence was detected that normalized replicate means did not follow a symmetric distribution. These findings were deemed as acceptable for using the unweighted OLS regression analysis.

#### 2.2.7 BMC and its Uncertainty

In the standard protocol the BMC was estimated directly from the best fit model. We also considered model averaging as an alternative option where, similar to the previous best fitting method, a number of suitable concentration–response models were fitted to the same data but in this case all resulting model fits were combined to provide an weighted average BMC estimates (Ritz, Gerhard and Hothorn, 2013). Uncertainty was always expressed as α=5%, i.e. the lower limit (BLL) corresponds to the 2.5% limit and upper limit (BUL) to the 97.5% limit. BLL and BUL were derived by three different methods, i.e. inverse regression, the delta method and bootstrapping. The estimation of the BMC and its 95% CI by model averaging was always performed in combination with bootstrapping. Inverse regression estimates both BLL and BUL directly from the regression fit around the BMC (Buckley, Piegorsch & West, 2009; Fang, Piegorsch & Barnes, 2015) and therefore puts high emphasizes on a successful regression fit in terms of robustness and reliability. The delta method is an asymptotic approach which combines information of the estimated model parameters to derive a Wald-type interval (Jensen et al. 2020). Bootstrapping uses computer-intensive simulation techniques that resamples the original dataset to create a huge number of so-called bootstrap samples, with each sample mirroring the original data set with an identical experimental design but newly simulated effect responses. On each bootstrap sample the same statistical data analysis was performed, resulting into a distribution of resampled BMC values around the original BMC estimation. If the median of this distribution equals the original BMC (unbiased resampling), then the 2.5% and 97.5% quantiles are expected to mirror the BUL and BLL of the original BMC, respectively. For each bootstrap sample, always the same regression model was used as part of the best-fit method, or one model-averaged BMC if model averaging was performed. To simplify the model averaging method, only three regression models were considered (four-parameter loglogistic, four-parameter Weibull and three-parameter exponential model). Bootstrapping was always conducted on 1000 resampled datasets, and due to the small sample sizes, we used always the parametric version (Efron, Bradley and Tibshirani, 1993). All resampling was performed by the function *bmdMA* of the R package *bmd* (Jensen *et al*., 2020). Bootstrapping can simulate a bootstrap sample which do not allow a BMC estimation or which leads to an unreliable BMC estimation that is well outside the tested concentration range. Therefore, a resampled BMC was excluded from the resampling distribution if it was 1.5-times above the highest test concentration or below the lowest tested concentration.

#### 2.2.8 Hazard Classification

CRSTATS uses a hazard classification approach which judges if data evidence is sufficient to define a compound as active for the specific DNT endpoint and if this can be distinguished from an activity observed in cell health related endpoints (viability and cytotoxicity). Accordingly, the endpoint-specific hazard of a compound is classified into five categories:

- **No hit**: no observed effect on the DNT-specific endpoint or on general cell health.
- **Unspecific hit**: the effect on the DNT-specific endpoint cannot be separated from an effect on the cell health related endpoint.
- **Borderline hit**: the separation between the effects on the DNT-specific endpoint and the effect on cell health related endpoint is statistically not clear (Leontaridou et al, 2017).
- **Specific hit**: the effect on the DNT-specific endpoint is clearly separated from an effect on the cell health related endpoint.
- **Not identified**: data are incomplete und do not allow any classification.

If the automatic classification failed due to a high uncertainty of the BMC or a missing BMC for the cell health related endpoint, the classification was recorded as **expert judgement** and classification into one of these five categories was done by manual inspection on the basis of all data evidence. An overview over all flagging alerts leading to expert judgement are given in Table S4.

The hazard classification approach was operationalized by hazard decision trees which reflect specific assay features and the directionality of the observed concentration response pattern (i.e. either reduction or inhibition). Common to all decisions trees is that they compare the BMC of the DNT-specific endpoints to the respective BMC of the unspecific endpoint (i.e., cytotoxicity or cell viability). For the NPC and UKN assays slightly different versions were developed, with all NPC assay endpoints accounting directly for the statistical uncertainties of both BMC estimations by using their corresponding CIs, and all UKN assay endpoints using pe-defined acceptance ranges instead. The principles of the hazard decision tree for data sets with decreasing concentration-response pattern (reduction) measured in NPC assays (NPC1, NPC2, NPC3 and NPC5, Table 1) are shown in Figure 3, and for increasing concentration-response pattern (induction) in Figure 4. Inductions are handled separately, because the specific and unspecific endpoints do not have the same relationship during an induction, compared to a reduction in the endpoint. A loss in general cell viability for example will likely result in an effect in cell proliferation, while an induction in cell viability does not necessarily increase cell proliferation. If migration (NPC2a) is affected, only cytotoxicity is used as a reference for all specific endpoints of NPC2-5. A reduction in migration also reduces cell viability due to the lower number of cells in the migration area and not necessarily due to cell death. If so, it cannot be used as valid reference to discriminate between a specific and unspecific effect. The same applies to effects in cell viability. In these cases, only cytotoxicity is used as general cell health reference for according specific NPC endpoints. More details can be found in the supplementary material (S1.3) and in table S3, and details about the classification tree applied to data from the UKN assays can be found in Masjosthusmann et al. (2020).

**Figure 3:**
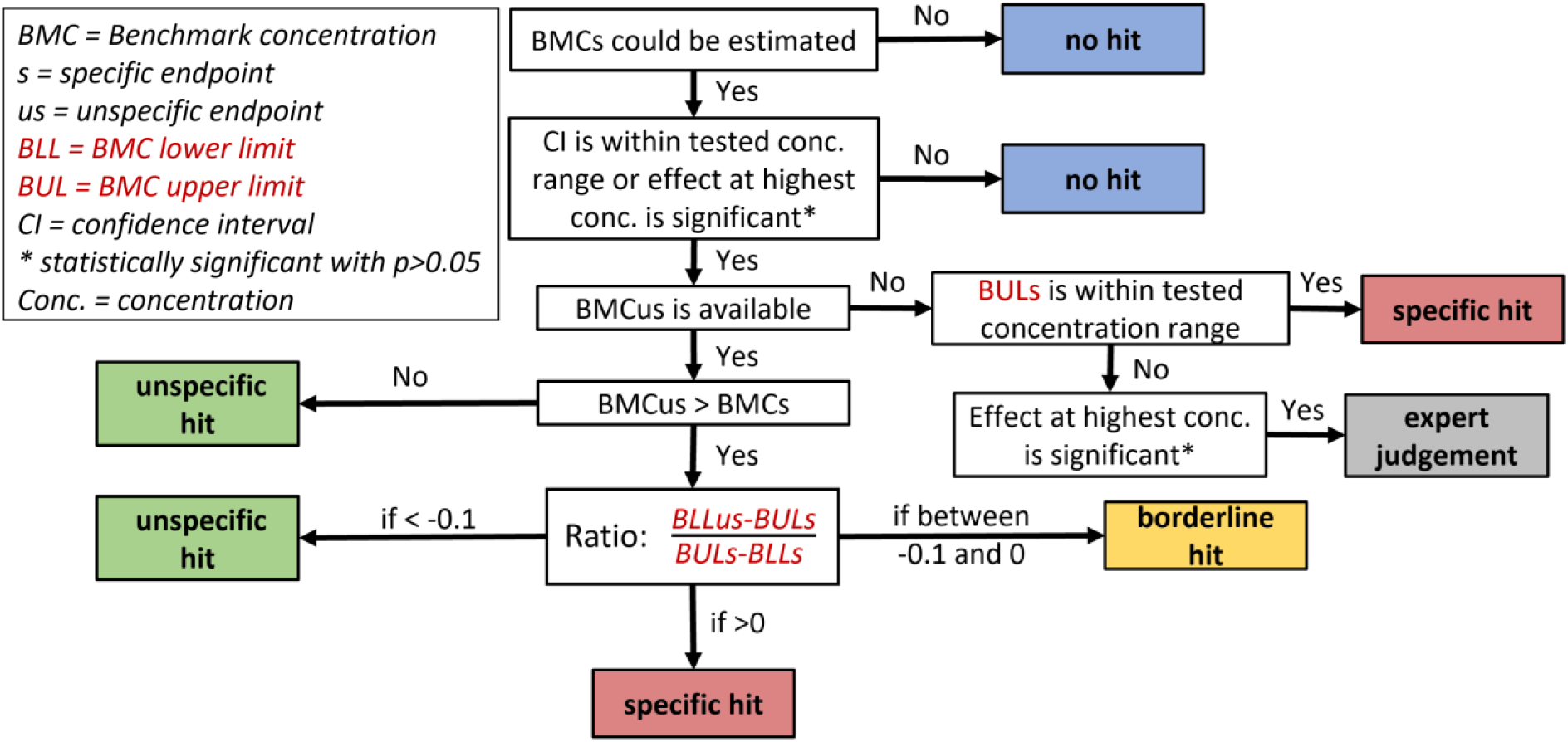
Decision tree for the NPC hazard classification of inhibitory effects. The decision tree shows for NPC1-5 data with decreasing concentration-response pattern how BMC estimations and their uncertainty (expressed as 95% confidence intervals, CI) for data from both specific and unspecific endpoints are used to classify the compound into one of the DNT hit categories (coloured boxes). Hits with the category “expert judgement” (grey box) will be classified into one of the DNT hit categories by manual inspection on the basis of all data evidence.

**Figure 4:**
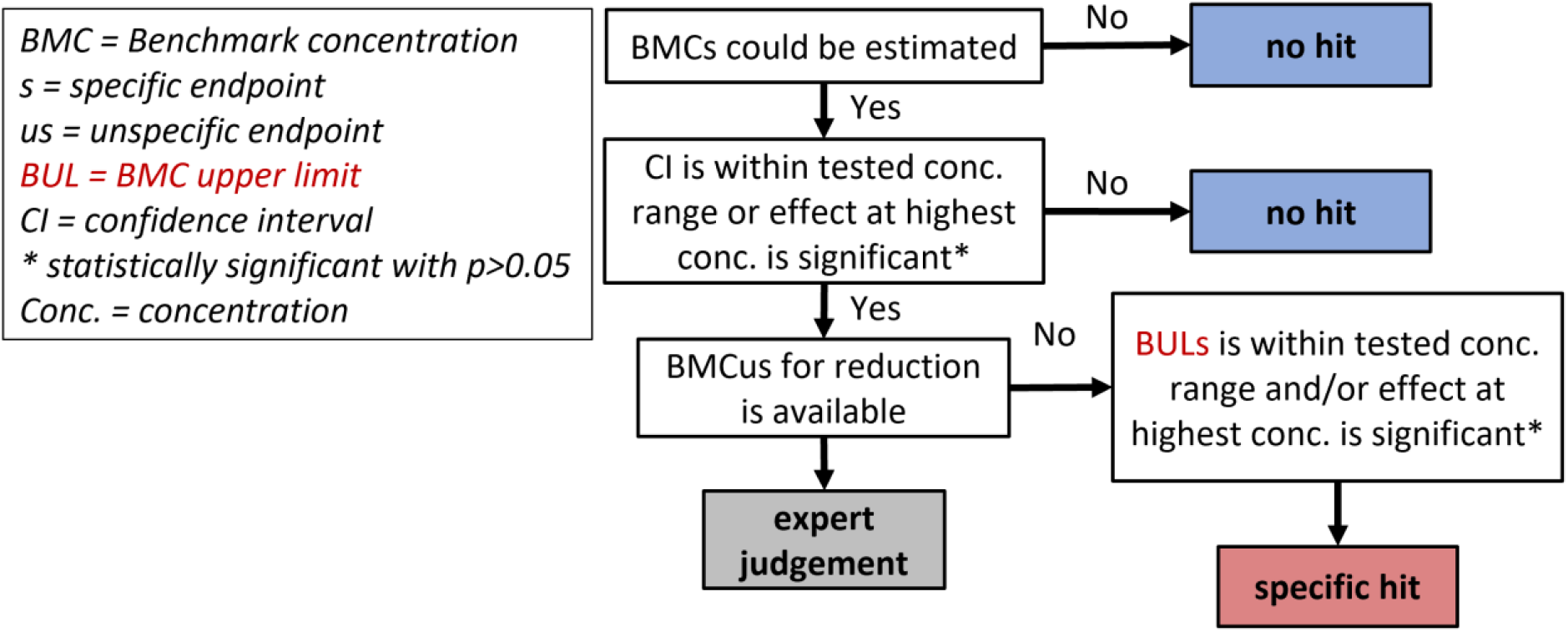
Decision tree for NPC hazard classification of increasing effects. The decision tree shows for NPC1-5 data with increasing concentration-response pattern (“induction”) how BMC estimations and their uncertainty (expressed as 95% confidence intervals, CI) for data from both specific and unspecific endpoints are used to classify the compound into a specific or no hit (coloured boxes). The presence of a cytotoxic responses can lead to an artefact in the DNT-specific endpoint and is therefore initially categorized as “expert judgement”. These hits will be classified into one of the DNT hit categories by manual inspection on the basis of all data evidence.

### 2.3 Assay Performance

From the 148 compounds tested in the DNT IVB, a set of 45 reference compounds (17 negative compounds that are known not to cause DNT; 28 positive compounds with proven DNT adversity in humans or mammals) was used for an evaluation of the DNY IVB predictivity. Hit decisions were derived from the hazard decision trees developed in 2.2.8, and the following performance parameters were used:

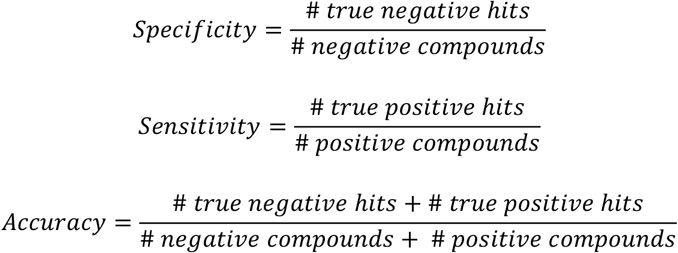

A negative compound was considered as true negative if it was not classified as specific hit or borderline in any of the assays. A positive compound was considered as true positive, if it was classified as specific hit or borderline in at least one assay.

## 3 Results

To perform a robust, fast and automated hazard characterization based on high content *in vitro* toxicity testing data, we have set up the R-based data evaluation pipeline CRSTATS (github.com/ArifDoenmez/CRStats), which offers multiple statistical options for the data evaluation of continuous concentration-response data. Based on these options, we have defined a standard data evaluation protocol (“standard protocol”) for DNT IVB data (Fig. 2), and by changing statistical methods as part of the protocol we studied their impact on the BMC estimation of the DNT IVB outcomes and the subsequent consequences for the hazard classification and overall DNT IVB performance (“alternative protocol”). The following statistical methods were chosen as alternatives to the standard protocol: 1) average replicate per experiment estimated by the arithmetic mean, 2) control normalization without re-normalization, 3) using a three-parameter log-logistic regression model (LL3rm) for the BMC estimation, 4) using model-averaging for the BMC estimation, 5) CI estimation of the BMC by the delta method, 6) CI estimation of the BMC by bootstrapping, 7) CI estimation of the BMC by model averaging, and 8) increasing the endpoint specific BMRs by 20%. Differences in the BMC estimation, the uncertainty of a BMC (expressed as the width of the central 95% confidence interval of a BMC estimation), the endpoint-specific hazard classification of the compound and the final assay performance were quantified and compared across the various specific assay endpoints.

In total, 148 compounds were tested on up to 14 DNT-specific and 8 cytotoxicity and viability endpoints, of which 2385 data sets fulfilled the minimal data requirements of the data evaluation pipeline. According to the standard protocol, it was possible to perform a regression analysis for 2385 data sets (1953 NPC and 432 UKN) and a hazard hit categorization for 1563 data sets from DNT-specific endpoints (1347 NPC and 216 UKN). In nearly one third of all best-fit model decisions the simplest regression model was chosen, i.e. the exponential function with two model parameters, followed by three-parametric models (55.7%) and by four-parametric models (10.3%). Only in 1.9% of all best-fit model decisions sufficient data were available to support the most complex regression model (5-parameter general log-logistic).

### 3.1 Impact of different data evaluation methods on the BMC estimation

To allow a better comparison of BMCs from different data scenarios, the BMC was transformed to a relative BMC on a log10 scale by relating the 100-fold BMC estimation to the highest test concentration of the data set:

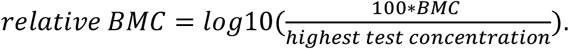

A relative BMC of 1 corresponds to a BMC that is tenfold below the highest test concentration of the data set, and a relative BMC above 2 corresponds to a BMC that has been extrapolated beyond the highest test concentration. The lower the relative BMC value, the more likely the estimation is supported by effect data from more concentrations.

The relative BMCs from the standard and alternative statistical protocols are shown in Figures 5 A-E for five statistical parameters that were changed, which the BMC of the alternative protocol always referring to the x-axis and the BMC of the standard protocol to the y axis. If a regression analysis could be performed but a BMC not established due to missing data support for the BMR, the BMC was flagged as “BMRnr” (BMR not reached) and included in the plot at the end of the BMR axes, i.e. a BMRnr value on the right side of the plot indicate a BMC estimation which was only possible for the standard protocol, and similarly, a BMC value on the top of the plot area indicate a BMC estimation that could only be established for the alternative protocol. Data sets for which none of the protocols were able to produce a BMC were excluded. Color-coded symbols refer to the 22 bioassay endpoints, and a data point on (or close to) the solid 45-degree line indicates a perfect agreement between the BMCs from both protocols. Three-fold BMC differences are highlighted by a belt around the line of perfect agreement (i.e. values outside of the belt have above three-fold change), and the percentage number of successful regression fits for the alternative protocol are included on top of each plot, with reference to the 1953 data sets for which a successful regression modelling was conducted according to the standard protocol. To identify general deviation patterns, we performed trend regression analyses between the relative BMCs, and the corresponding value of the goodness-of-fit criterion (R^2^) is provided in the plot: the higher the coefficient, the more consistent the results between the two protocols. For the trend analysis, we set a relative BMC = 2.47 for a BMRnr, i.e. a 3-fold difference between the highest concentration and a fictional BMC was assumed. Not shown are BMC differences for the bootstrapping and delta method, as both refer to the same BMC and thus would have resulted always into identical BMCs in the plot.

**Figure 5:**
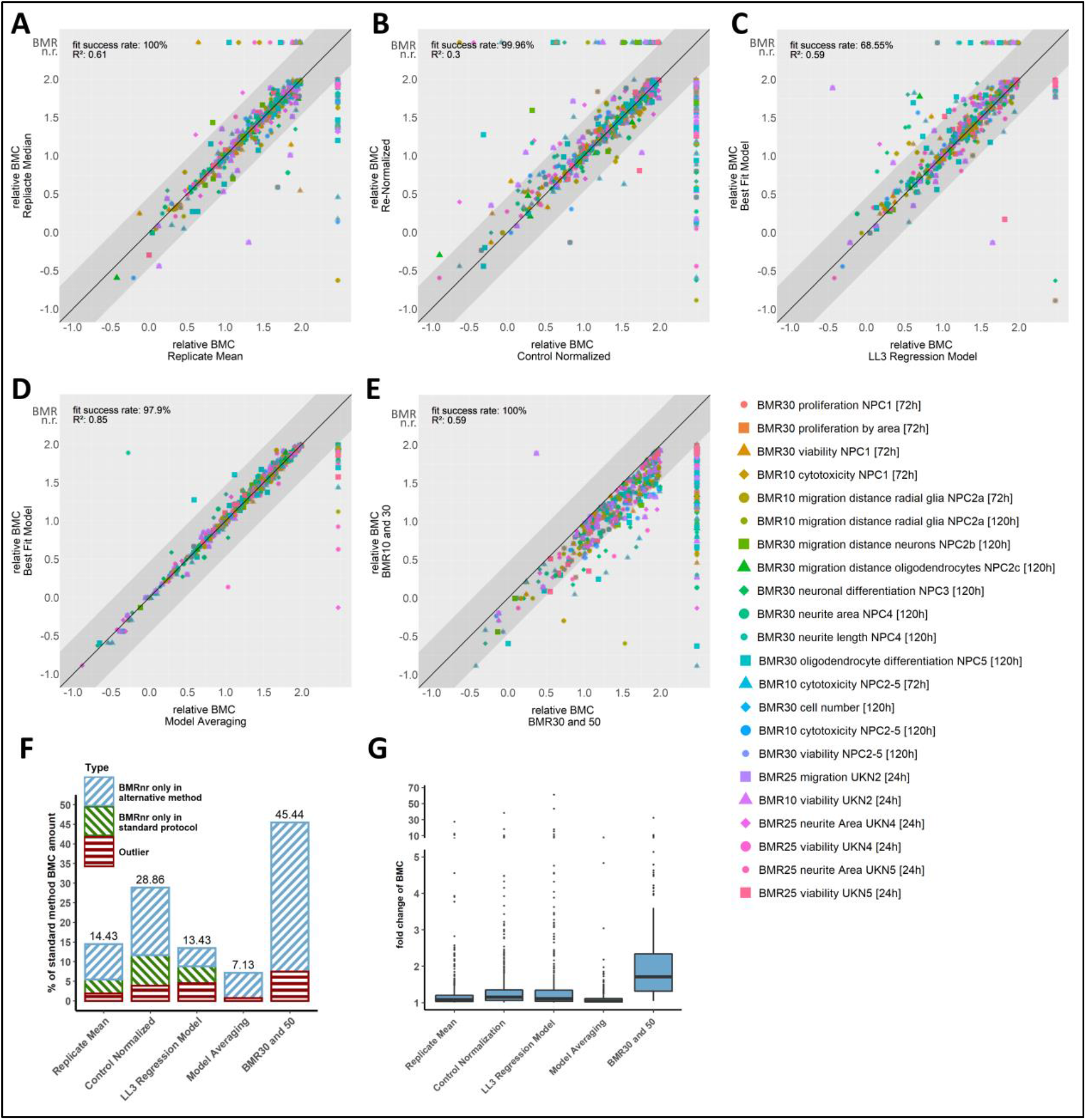
Impact of methodological changes in the data evaluation on the BMC estimation. BMCs for 148 compounds tested on up to 22 endpoints from 8 assays were estimated using the standard protocol and opposing alternative methods. A-E): A relative BMC was expressed as the log10-transformed ratio between the 100-fold BMC and the maximum test concentration, and relative BMCs from all data sets and endpoints but different statistical methods were plotted against each other. The solid black trend line indicates no differences between the relative BMCs, the grey interval around the trend line indicates values within a three-fold range. Values outside this interval are considered as relevantly different between the opposing methods. If a relative BMC could be calculated for only one method, the missing value of the opposing method is plotted as BMRnr area on the right or upper side of the graph. Relative BMCs are colored according to their bioassay endpoint. To indicate the strength of agreement between both data evaluation protocol, the goodness-of-fit coefficient from a trend regression analysis between both relative BMCs is included (R^2^; top left), and the percentage of successfully applied regression models of the alternative protocol in relation to the standard protocol is shown top left (“fit success rate”). A) Experimental median replicates versus mean replicates (n = 568). B) Re-normalized data versus control-normalized data (n = 630). C) Best fit approach versus a predefined three parameter log-logistic regression model (n = 520). D) Inverse regression versus model averaging (n = 604). E) BMR10+30 (BMR10+25 for UKN) versus BMR30+50 (n = 604). F) Percentage of all data sets for which the protocol change lead to a BMC change in terms of BMRnr (i.e. a BMC could not be determined from the regression fit) or an above three-fold BMC change. G) Distribution of BMC fold-changes in response to statistical method changes from the standard protocol. Box whisker plot shows the median (horizontal line), interquartile range (box), 5% and 95% percentile values (whisker), and extreme values (black dots).

We found the most profound BMC differences between the data re-normalization and control normalization (Fig. 5B), with an R^2^ of 0.3. The main reason for the huge number of BMC disagreements is due to huge number of BMRnr’s, i.e. regression fits that could establish a reliable BMC for the endpoint-specific BMR in only one of the protocols. Using the mean as replicate average instead of the median (Fig. 5A), using a predefined regression model (LL3rm) instead of the best fit method (Fig. 5C), and using a higher BMR resulted in moderate BMC changes, with R^2^’s between 0.59-0.61. The best agreement between relative BMC values was observed for the comparison between the outcomes from model averaging against the best-fit method (Fig. 5D) with an R^2^ of 0.85.

The number of datasets for which a regression model could be fitted for the alternative protocol was related to the number of fits for the standard protocol and expressed as relative “fit success rate”. All changes of statistical methods lead to similar success rates, with the exception of the sole application of the three-parameter log-logistic model which led to a noteworthy loss of successful regression fits (68.55% success rate).

To further explore differences between BMC estimates, the number of BMRnr cases that only occurred in the alternative protocol (i.e. the standard protocol did result in a BMC while the alternative protocol did not; Fig. 5F, blue shaded area of bar), the number of BMRnr cases that only turned out in the standard protocol (Fig. 5F, green shaded area of bar) and large differences outside the belt (“outliers”, Fig. 5F red shaded area of bar) were compared to the total number of BMCs that were estimated by the standard protocol. Most protocols that lead to less successful BMCs were caused by the inability of the data to support the regression modelling for the intended BMR level. All alternative protocols together led to less BMCs but more BMRnr cases, with protocol changes to control normalization and higher BMRs resulting into the highest increase towards BMnr cases (i.e. less BMCs), with an increase of 17.25% and 37.98% of BMRnr cases, respectively. Taking only the cases with huge BMC differences into account (“outliers”), the number of BMCs that were either lost or gained due to the protocol change was further quantified: model averaging led to the smallest number of relevant changes (7.13%), followed by replicate averaging by mean, fixed regression model (LL3rm) and control-normalization with moderate changes (13.43%-28.86%), up to >40% changes were reached if a higher BMR was used.

Differences between the relative BMCs were also expressed as fold-change, and the distribution of all fold changes summarized as median and the interquartile ranges (IQR) (Figure 5G). Here, the BMRnr cases were excluded from the fold change analysis. In alignment to the previous results, the protocol change towards higher BMRs led to the most severe fold-change (median = 1.71, IQR = 1.02). The protocols with mean replicate average, control-normalized data, model averaging and choice of higher BMRs showed moderate median fold-changes of estimated BMCs ranging from 1.04-1.15 (IQR ranging from 0.09-0.3).

### 3.2 Impact of different data evaluation methods on the BMC uncertainty

Next, we analyzed how changes to the standard evaluation protocol can influence the overall uncertainty of the BMC estimation. The uncertainty of a BMC is estimated as central 95% confidence interval with the BLL corresponding to the lower 2.5% interval section and the BUL to the upper 97.5% interval section. The width of the interval (i.e. the difference between the BUL and the BLL) is an essential factor in some of the classification models of the hazard characterization. Similar to the analysis of BMC changes, a CI width was transformed to a relative CI width by fixing it to the maximal test concentration of the data set:

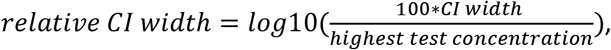

with CI width = BUL – BLL, where BLL and BUL are the 2.5% and 97.5% confidence interval of the BMC estimation. A single relative CI width has no meaningful interpretation, and it was only used in combination with a second value with reference to the same highest test concentration.

Changes to the standard protocol which lead to changes in the relative CI width were visualized in the same way as in the previous section, with the relative CI widths from the standard and alternative protocol shown as endpoint-specific symbols for all data sets in a common scatter plot, with each plot referring to a specific method change (Fig. 6A-G). Values below the line of perfect agreement indicate an increase of the CI width, i.e. the BMC estimation of the alternative protocol is considered as more uncertain, and values above the line indicate a more certain BMC estimation than judged by the standard protocol. Also, a supporting trend analysis was conducted, with the corresponding goodness-of-fit criterion (R^2^) provided in the plot, and a belt around the line of perfect agreement between both relative CI widths was included, with larger than three-fold changes outside this belt considered as relevant.

**Figure 6:**
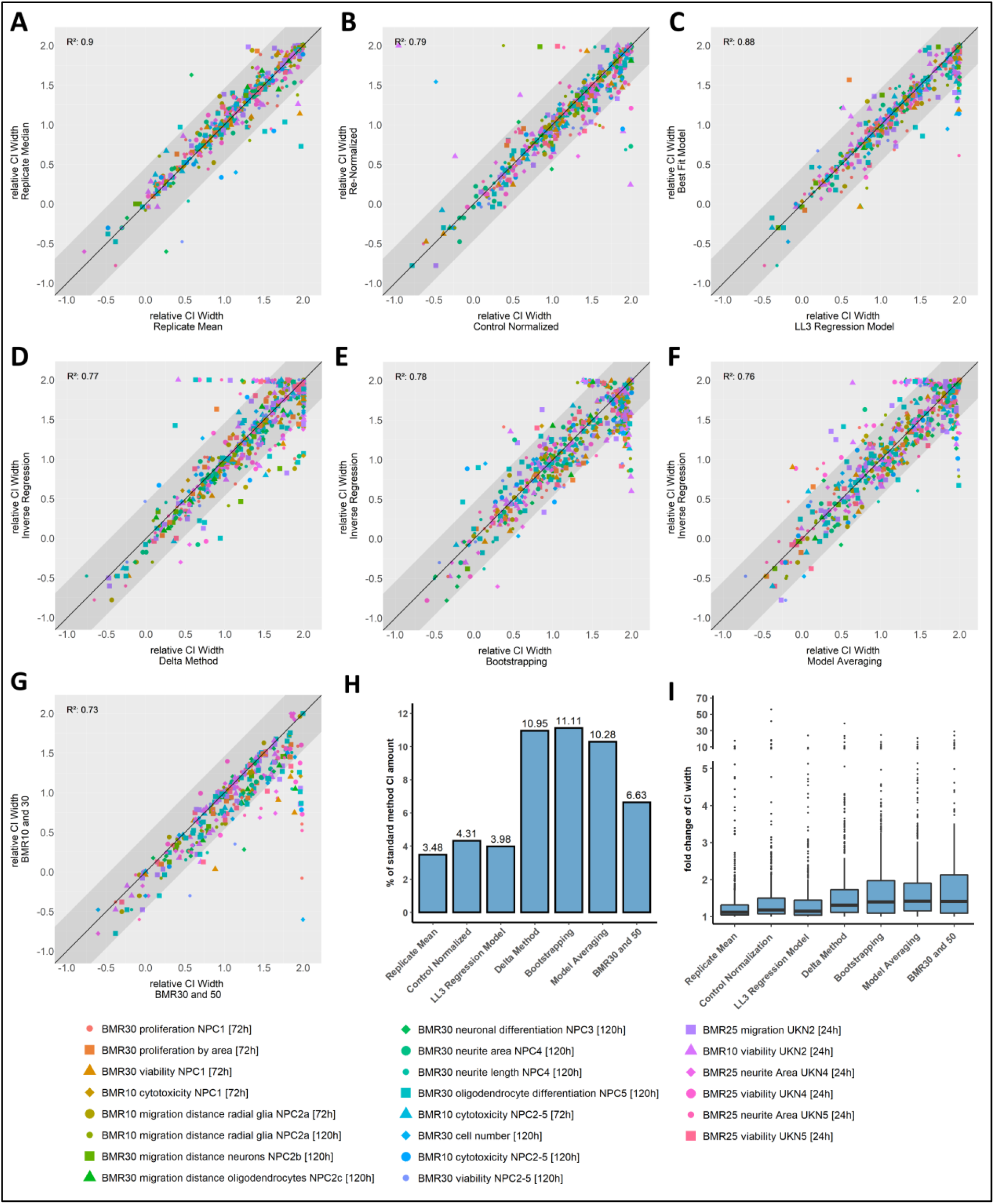
Impact of methodological changes in the data evaluation on the uncertainty of a BMC estimation. The BMC uncertainty was expressed as the width of the central 95% confidence interval (CI) around the BMC estimation, and CI widths for 148 compounds tested on up to 22 endpoints from 8 assays were determined using the standard protocol and opposing alternative methods. A-G): a relative CI width was calculated as the log10-transformed ratio between a 100-fold CI width and the highest test concentration, the relative CI width of the alternative protocol was plotted against the relative CI width of the standard protocol. The solid black trend line indicates identical CI width’s, the grey interval around the trend line indicates values below three-fold change. Values outside of this interval are considered as relevantly different between the opposing methods. Endpoints are indicated by a different color and shape. The goodness-of-fit coefficient from a trend regression analysis between both relative BMCs was calculated (R^2^; top left), indicating the strength of agreement between both data evaluation protocols. A) Experimental median replicates versus mean replicates (n = 517). B) Re-normalized data versus control-normalized data (n = 502). C) Best fit approach versus a predefined three parameter log-logistic regression model (n = 499). D) Inverse regression versus delta method (n = 588). E) Inverse regression versus bootstrapping (n = 600). F) Inverse regression versus model averaging (n = 561). G) BMR10+30 (BMR10+25 for UKN) versus BMR30+50 (n = 359). H) Percentage of all BMC for which the protocol change lead to a threefold change in the relative CI in relation to the total number of BMC estimations. I) Distribution of CI width fold changes in response to statistical method changes from the standard protocol. Box whisker plot shows the median (horizontal line), interquartile range (box), 5% and 95% percentile values (whisker), and extreme values (black dots).

Outcomes of the trend analyses show that protocol changes due to the experimental mean replicate or the sole application of the LL3rm regression model led to the least impact on the CI width, with R^2^s of 0.9 and 0.88, respectively (Fig. 6A and 6C). All remaining protocol changes led to slightly higher changes in the BMC uncertainty, with R^2^ between 0.79-0.73 (Fig. 6A-G). The number of increased or decreased confidence intervals around the BMC was balanced across all protocol changes, with the exception of the protocol change towards higher BMRs where the BMC uncertainty was increased for the majority of data sets. The total number of BMC estimations for which a method change led to changes in the CI width that we consider as relevant (i.e. values outside belt around the perfect agreement) was then compared to the total number of BMCs: all method changes directly involved in the estimation of the BMC uncertainty (delta method, bootstrapping and model averaging) changed the CI width of the BMC for ca. 10% of all BMCs, whereat method changes that are expected to impact the uncertainty estimation of a BMC only indirectly (mean replicate average, control-normalized data, sole application of an LL3rm, higher BMRs) had a minor impact on the determination of the BMC uncertainty (Fig. 6H). The distribution of fold change of CI widths is shown in Figure 6I.

### 3.3 Examples

In the following we have selected five data examples which demonstrates the impact of methodological changes in the standard data evaluation protocol on a BMC estimation (Figure 7).

**Figure 7:**
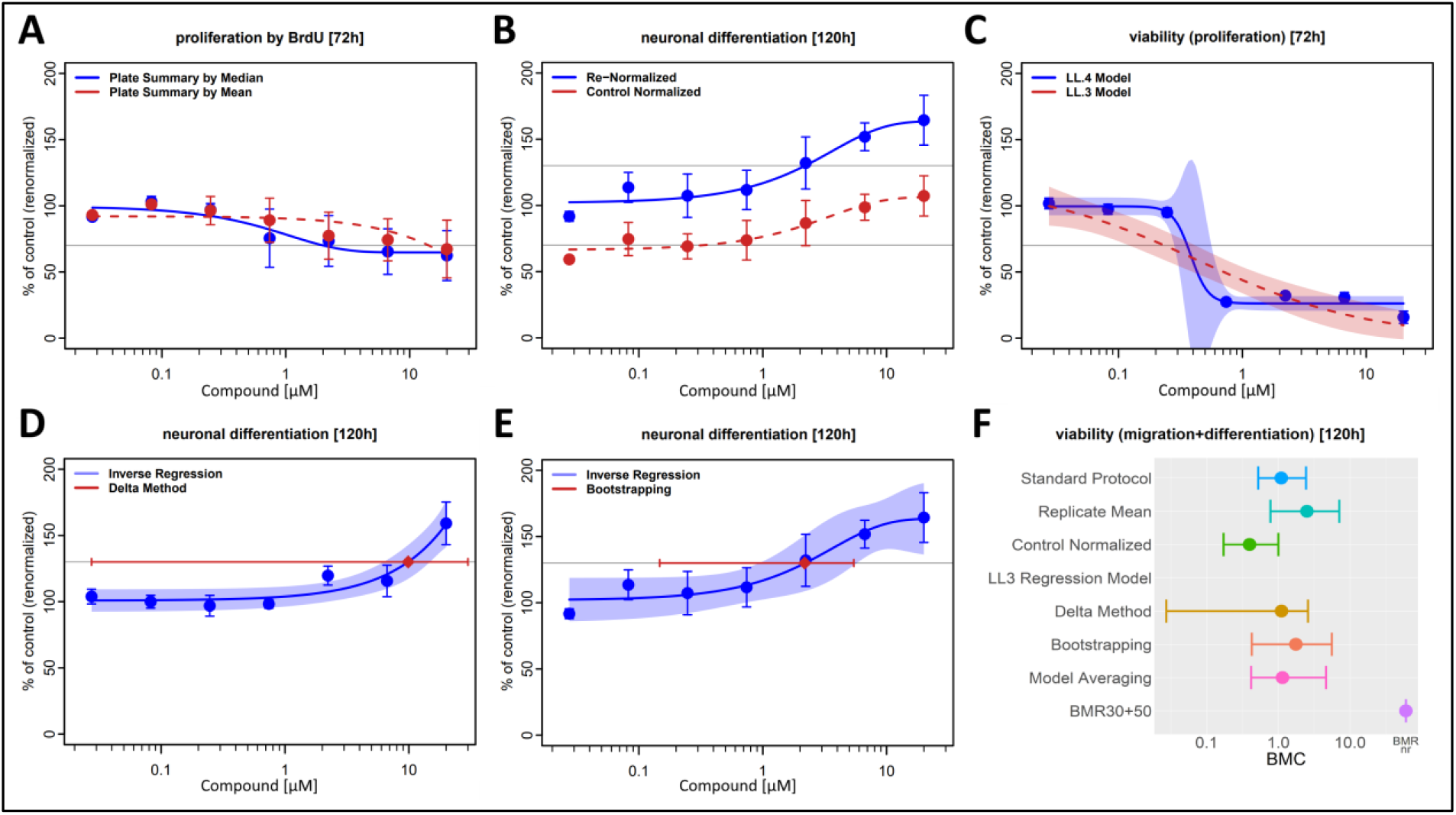
Data set examples and the impact of methodological changes in the data evaluation on the regression modelling and the BMC estimation and its uncertainty. A)-E) For several different steps of the data analysis and evaluation, the data resulting from the standard protocol (blue) is compared to the data deriving from the alternative protocol (red). Error bars show the SEM between summarized experiment data. Horizontal grey lines indicate the BMR. A) Experiment summarization by median and by mean. B) Re-normalized data and control-normalized data. C) Best fit approach and use of only a LL3 regression model. CI is displayed as confidence band around the fit model. Both models are applied to the data shown in blue. D) Inverse regression and delta method. CI of the alternative method is shown as red bar and BMC as red square. E) Inverse regression and bootstrapping. F) All method changes and their resulting BMC (displayed as dots) and CI values (displayed as bars) are shown for one exemplary dataset.

i. Replicate median versus replicate mean (Figure 7A, proliferation by BrdU after 72h exposure): response data were calculated either as the median (blue, standard protocol) or as the arithmetic mean (red, alternative protocol) of the replicate responses and expressed as mean ± SEM (n=5). The corresponding best-fit regression models are shown as solid (standard protocol) and dashed lines (alternative protocol), with the horizontal line corresponding to a BMR30. The combination of a large data variability and presence of individual data outlier led to mean response estimations closer to the control level, which was reflected by distinct regression curve estimates for both protocols. As consequence, a 7-fold higher BMC30 value was estimated for the alternative protocol, with a BMC30 of 2.02 μM according to the standard protocol and a 14.3 μM for the alternative protocol.
ii. Re-normalization versus control normalization (Figure 7B, neuronal differentiation after 120h exposure): response data were either normalized to the average response observed at the lowest test concentration (blue, standard protocol) or to the controls (red, alternative protocol). The four lowest test concentrations produced similar control-normalized responses between 60-70%, with no indications for a trend between them. A regression analysis on all control-normalized responses suggests that only the highest test concentration produced a response not distinguishable from the controls, which we deemed as unrealistic, and, as consequence, the data set was not considered for a reliable BMC estimation. However, data re-normalization led to a more valid induction pattern, and the BMC estimate from the best-fit regression model was accepted.
iii. Best-fit model selection versus a pre-defined regression model (Figure 7C, cell viability after 72h exposure): re-normalized response data are presented in the same way as in the previous examples, with the blue regression curve corresponding to the best-fit regression model (standard protocol) and the red curve to the three-parameter log-logistic model (alternative protocol). In this example, the four-parameter log-logistic model (Table S2) was selected as best-fit regression model. In addition, the 95% confidence intervals around the entire curve estimates are included for both regression models. The exposure concentrations produced either no or maximal responses, with no data support for effect responses between. In a strict statistical sense, this data pattern does not allow the estimation of a reliable data curve, which is indicated for the three-parameter model (alternative protocol) by its poor data description and for the four-parameter log-logistic model (standard protocol) by its huge confidence belt for intermediate effect estimates. Nevertheless, the BMC30 (and its uncertainty) derived from the four-parameter log-logistic model (0.347 μM, 95% CI: 0.272-0.572 μM) seems reasonable and it is unlikely that a re-testing of the same compound on a refined concentration range will contradict this BMC30. This example demonstrates that although a BMC estimation might not fulfill all criteria according to “best statistical practice” (and be judged as unreliable by statisticians), it still can provide sufficient information to be assessed by the experimenter as reliable.
iv. Uncertainty estimation of the BMC by inverse regression versus the delta method (Figure 7D, neuronal differentiation after 120h exposure): the 95% confidence intervals of the BMC30 estimated from the same best-fit regression model and data set are shown either by inverse regression, i.e. the interval along the horizontal 130% response line that intersects with the confidence belt of the regression curve (blue, standard protocol), or by the delta method (red). The confidence belt of the BMC30 by inverse regression provides a reliable expectation about where a BMC30 can be expected if the same experiments would be repeated, whereat the delta method provides a confidence belt which spans the entire test concentration range and therefore provides the misleading conclusion about a non-existing data support.

(iv) Uncertainty estimation of the BMC by inverse regression versus bootstrapping (Figure 7E, neuronal differentiation after 120h exposure): similarly to the previous example, 95% confidence intervals of the BMC30 estimated from the same best-fit regression model and data set are shown for two different statistical methods, and similar to the delta method, bootstrapping provides a large 95% confidence belt of the BMC30 which could be interpreted misleadingly as lacking data support for the regression modelling. The most likely reason for the poor performance of the bootstrap is the combination of a relatively large data variability and the responses observed at the second lowest concentrations which are not well described by the regression model.

Finally, we selected a single data set and analyzed it according to the standard protocol and all the methodological changes we have conducted to estimate a BMC and its 95% CI (cell viability, Figure 7F). It summaries well how the different statistical methods can change a BMC estimation: a control-normalization of the response data shifted the BMC and its CI to much lower test concentrations, application of the delta method led to a drastic increase of the CI of the BMC30, the sole application of the LL3rm regression model failed and did not provide any estimation and increasing the BMR30 to a BMR50 led to a non-estimable BMC (BMRnr).

### 3.4 Method impact on hazard classification

An important application of the BMC estimation is the endpoint-specific hazard classification of the test compound into one of five hit categories, i.e. if the compound produced sufficient data evidence to be judged as a DNT-specific hit, borderline hit, unspecific hit, no hit, or as not identifiable (due to missing data support). Although all decision trees were setup as automatic systems, some data scenario provided insufficient data and were flagged for an expert judgement. The number of data scenarios for which the hazard classification was performed by “expert judgement” are listed in Table 2 for the standard protocol and seven methodological changes, divided according to the main decision trees developed for data from NPC or UKN assays. In total, 1563 classifications were conducted (NPC: 1347, UKN: 216), of which 68 (NPC: 30, UKN: 38) were flagged for an expert judgement according to the standard protocol. All protocol changes led to similar numbers, with the exception of the delta method applied to data outcomes from NPC assay endpoints which required expert input for three-times more classifications. A marked difference was observed between the decision trees for NPC and UKN assay endpoints, with up to 5 times more classifications flagged for expert judgement for UKN outcomes depending on the statistical method chosen.

**Table 2:** Number of endpoint-specific DNT hit classifications judged by experts. The numbers of hit classifications by expert judgement are presented as percentage of all classifications that were supervised by the hazard decision trees.

| Method | NPC [%]<sup>1</sup> | UKN [%]<sup>2</sup> |
| --- | --- | --- |
| Standard Protocol | 2.23 | 17.59 |
| Replicate Mean | 2.15 | 16.67 |
| Control-normalized | 3.27 | 15.74 |
| LL3rm | 2.23 | 12.50 |
| Delta Method | 8.09 | 18.52 |
| Bootstrapping | 3.12 | 17.13 |
| Model Averaging | 2.90 | 15.74 |
| BMR30+50 | 1.93 | 12.96 |

<sup>1</sup>NPC = Data outcomes from NPC assays, <sup>2</sup>UKN = Data outcomes from UKN assays

Due to the poor performance outcomes of the delta method in judging the uncertainty of a BMC estimation and the consequence of a more likely expert intervention in the automatic hazard classification, we judge this method as too unreliable and have excluded it from all remaining analyses.

Exemplary data sets are shown for three different classification scenarios: (i) a specific DNT hit decision for a significantly inhibited oligodendrocyte differentiation at exposure concentrations above 0.25 μM, but only a marginally reduced cell viability (marker for cytotoxicity) at 20 fold higher concentrations (Figure 8A), (ii) an unspecific hit decision for a significantly inhibited oligodendrocyte differentiation and cytotoxicity observed at same concentration ranges (0.24 to 2.2 μM) (Figure 8B), and (iii) a data scenario which was flagged for an expert judgement because the highest test concentration (20 μM) produced a weak but significant effect reduction for the specific endpoint but the regression analysis estimated a BMC10 (and BLL) that was outside the test concentration range. On closer inspection of the experimental data (Figure 8C, with each color-coded symbol representing the replicate median from an independent experiment) it was decided that responses from both the specific and unspecific endpoint were not distinguishable, and thus the weak response reduction of the specific endpoint was classified as unspecific.

**Figure 8:**
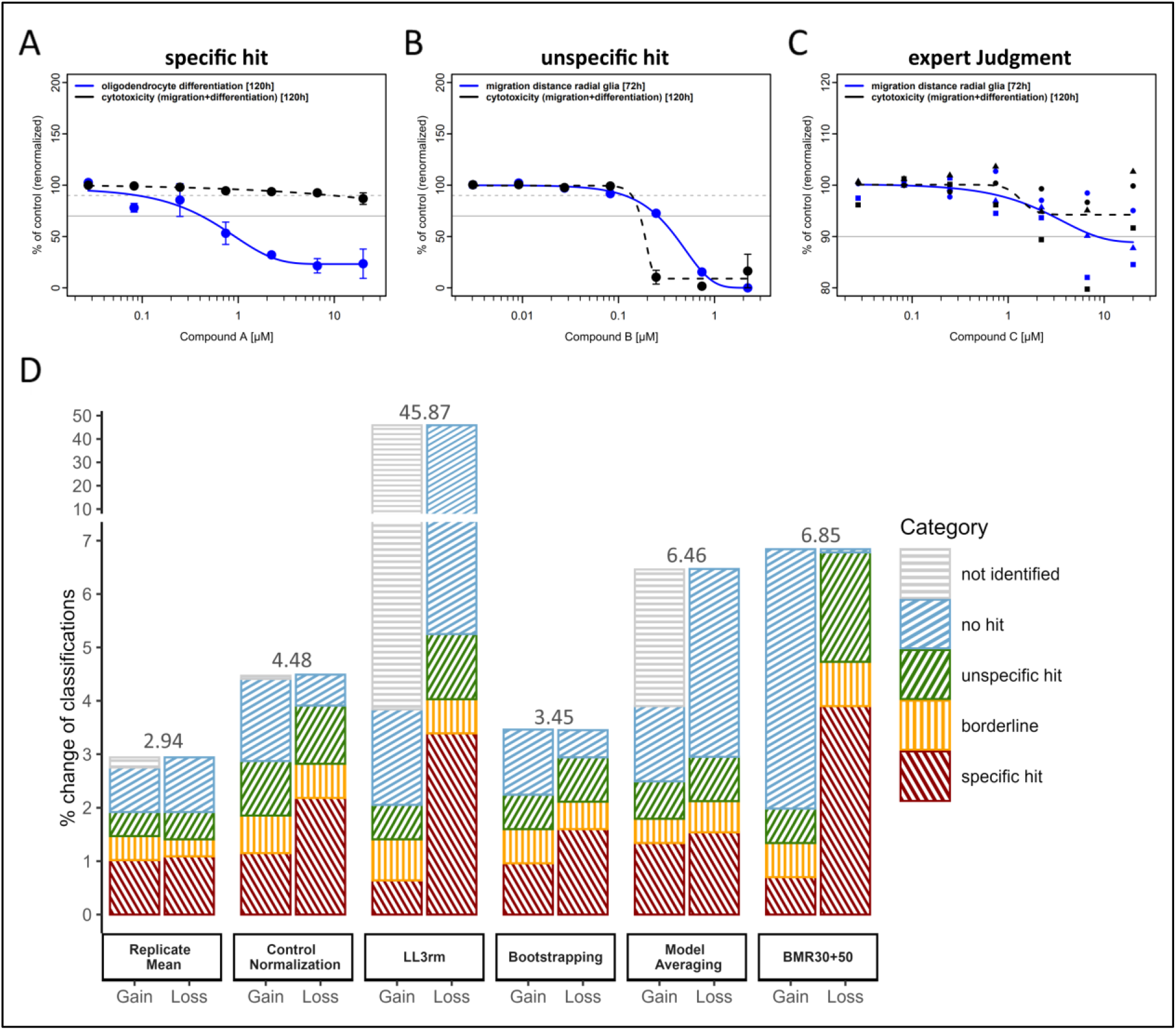
Number of endpoint-specific DNT hit classification changes in response to changes in the standard data evaluation protocol. A-C) Exemplary data sets for three different classification scenarios: concentration-response data from the specific (blue) and unspecific (black) endpoints are from 5 independent experiments, with effect responses re-normalized to the regression estimate at lowest test concentration and summarized as mean±SEM. Horizontal lines indicate the BMR levels for the BMC estimation, where straight lines indicate the specific endpoint BMR and dotted lines the unspecific endpoint BMR (if they differ). Data were always analyzed according to the standard data evaluation protocol A) Specific hit: the specific endpoint (oligodendrocyte differentiation) is impacted at non-toxic concentrations. B) Unspecific hit: inhibition of oligodendrocyte differentiation and cell viability are observed at similar concentration ranges. C) Hit classification by expert judgement: an automatic hit classification was prevented by ambiguous data, but judged as “unspecific” by experts. D) For each methodological change to the standard protocol, the number of hit changes is expressed as percentage of the total number of hit classifications, divided into in “gains” (i.e. the percentual increase of hazard hits in relation to the standard protocol) and “losses” (i.e. the percentual decrease of hazard hits in relation to the standard protocol). Different bar segments represent the different classification categories.

Figure 8D provides an overview about the total number of hit classifications that changed in response to changes of the standard protocol. Expressed as percentages and for each methodological change, the changes of hit classifications are further divided in “gains”, i.e. the percentual increase of hazard hits in relation to the standard protocol, and “losses”, i.e. the percentual decrease of hazard hits in relation to the standard protocol. Here a change toward replicate averaging by mean, control normalization and bootstrapping caused the lowest number of classification changes (<5%), followed by methodological changes towards model averaging or higher BMR levels which led to almost 7% different hit classifications. Here, model averaging increased the number of “not identified” classifications by 2.56%, mostly at the cost of “no hit” classifications, and the higher BMR levels led to 4.86% more “no hit” classifications (in line with the data example of Figure 7G). The by far most severe changes of hit classifications were observed if only the LL3rm regression model was used to describe the experimental concentration response data (45.87% total difference), which led to 42.03% more “not identified” classifications. The latter is most likely the consequence of unsuccessful regression modelling (and corresponding BMC estimation) due to lack of sufficient data support for this model (see 3.1 and 3.2).

### 3.5 Assay performance

To assess how changes in the data evaluation protocol might impact the evaluation of the DNT IVB’s predictivity, 28 reference chemicals of known DNT and 17 negative control chemicals were selected (Masjosthusmann et al. 2020), with all 45 substances tested in the DNT IVB, and the overall performance of the DNT IVB was quantified by its specificity, sensitivity and accuracy. Outcomes are shown for the standard protocol as well as all relevant changes in Figure 9: (i) Specificity (Fig. 9A): standard protocol and changes of it led always to a specificity between 87.5% and 100%, i.e. a truly DNT negative substances were almost always also judged as negative by the DNT IVB, and the standard protocol seems to be robust against methodological changes in judging false-negatives. (ii) Sensitivity (Fig. 9B): 23 of the 28 DNT substances (82.1 %) were successfully identified by the DNT IVB if the standard protocol was used, but changes to it led always to a lower sensitivity. (iii) Accuracy (Fig. 9C): The best performance was achieved for the standard protocol (88.6%), followed by a methodological change to bootstrapping (86.4%), higher BMR levels (84.1%), mean replicates, control normalization, pre-defined regression model (all 81.8%) and model averaging (77.3%). The latter performed 11.3% below the accuracy value of the standard protocol. A detailed overview over the hit definition of all control compounds is given in supplementary segment 2.1 (Tab. S5-S7).

**Figure 9:**
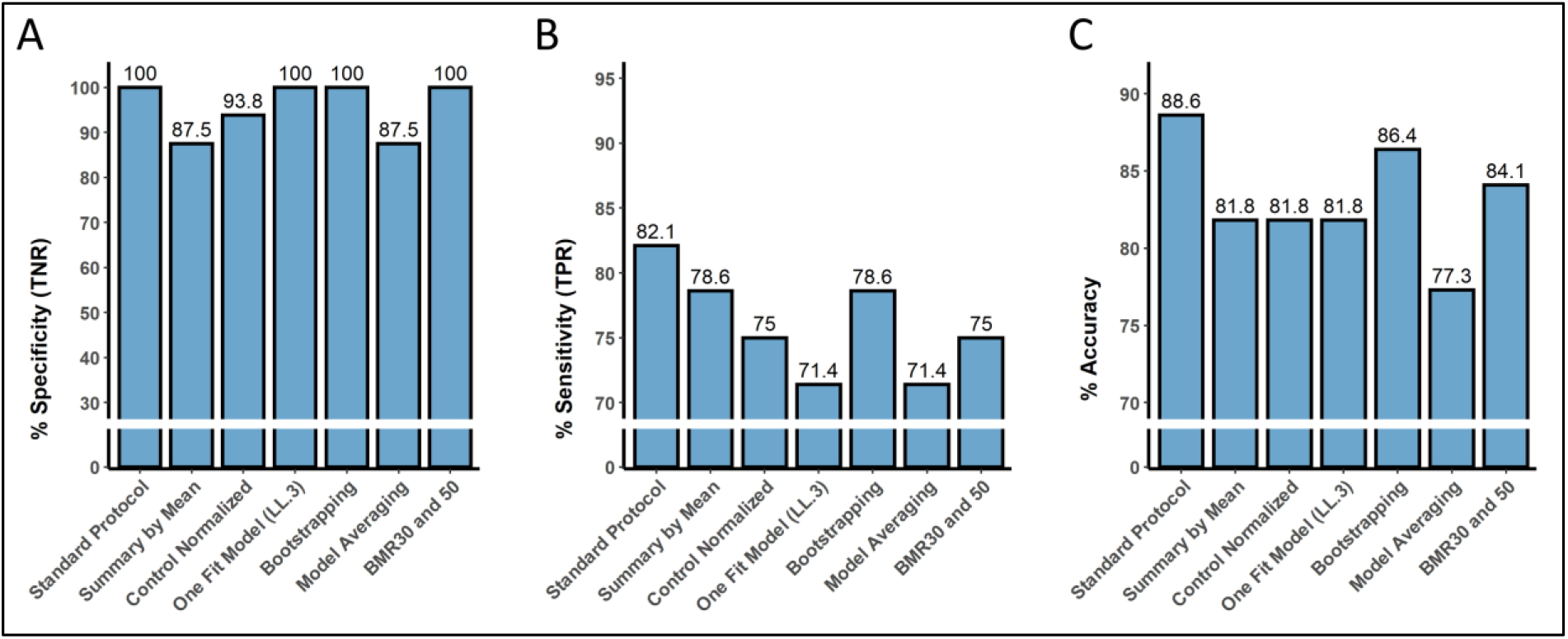
Evaluation of the predictive performance of the DNT IVB based on the standard data evaluation protocol and changes. Bar graphs show the results of the predictive capability of the DNT IVB for 28 substances of known DNT and 17 negative control substances in terms of specificity, sensitivity and accuracy.

## 4 Discussion

The basis for this biostatistical study is a compound screening project performed on behalf of an EFSA procurement during the years 2017-2020 (OC/EFSA/PRAS/2017/01). Twelve DNT test methods with accompanying cytotoxicity and viability assays belonging to an OECD DNT IVB (Crofton and Mundy, 2021) were challenged with 124 compounds from different compound classes including expected negative control compounds (Masjosthusmann et al. 2020). This paper is not about informing on the compounds’ effects on specific neurodevelopmental key events, which can be found elsewhere (Masjosthusmann et al. 2020; Blum et al. in revision), but rather analyzes the impact of common biostatistical concentration-response methods on the overall DNT IVB performance. As in vitro methods have been gaining complexity over the last decade, i.e. from simple reporter gene assays towards organotypic cultures, we tested the hypothesis if the selection of a biostatistical method can affect the performance of the DNT IVB. Specifically in the field of developmental toxicity, where in vitro test systems can nowadays assess biologically more complex systems, like changes in key developmental processes over time, such an evaluation seems timely. Hence, a comparative assessment of different biostatistical methods on the BMC estimation, DNT hit classification and DNT IVB performance was performed.

### 4.1 Experimental mean or median replicate

Instead of the individual experimental readouts, we used always their “average” response per test concentration and experiment (replicate average) as statistical unit in the concentration-response regression analysis. Our main argument for this data reduction was that the BMC and BLL estimation should reflect mainly biological and between-study variability, with the advantage that less complex statistical methods are required, which is a crucial requirement for the robustness of an automatic data evaluation pipeline. An “average” of the replicate responses can be estimated in various ways, with the arithmetic mean calculation the most popular and statistically often best option if the response data follow the rules of a symmetric distribution. However, the presence of an outlier can violate this assumption, and as a consequence it can lead to a biased estimation of an average that does not represent the observed data correctly. To protect the mean against an outlier would not only require outlier detection methods, which are per se problematic for small sample sizes, but also a decision on how to handle these values in further data analyses (e.g., removing, winsorization, trimming). A common alternative to the mean is the median which is more robust against outliers, but also known to be a more uncertain estimate for an average (Maindonald and Brown, 2010). As assay endpoints of the DNT IVB can produce a relatively high data variability within an experiment, we ruled out outlier detection methods and considered the median of the replicate responses as default option for the standard protocol.

Our study shows that the alternative of using the arithmetic mean led in average only to minor changes in BMC estimations and hazard classification outcomes, which might refer to those data sets where either no outliers were present or outliers occurred at concentrations with only little influence on the regression analysis. Nevertheless, for a few data sets a decision towards the median or mean had a strong influence on the best-fit regression analysis such that, at worst case, the subsequent BMC estimation was prevented (“BMRnr”). Although the choice of the average replicate calculation had in comparison to other methodological protocol changes only a minor impact on the hazard classification, it still lowered all performance parameters (specificity, sensitivity, accuracy) that we used to assess the DNT IVB’s predictive power for identifying DNT adversity. Altogether, our study outcomes strengthen the argument for using the median.

### 4.2 Data normalization to control and re-normalization

Typically, readouts from *in vitro* endpoints can substantially vary between experiments and thus require a normalization to make them comparable across experiments. The classical approach is to “anchor” all values to the average response of a negative control (and, depending on the assay endpoint, positive control). The expectation is that an exposure concentration with no impact on the assay endpoint will produce readouts similar to those from the control reference, which in the concentration response context means that a regression curve is expected to “equal” the control responses at zero and non-effective low exposure concentrations. This expectation does not always hold true in experimental practice, and readouts from concentrations that are expected to show no exposure activity (on basis of mechanistic reasoning or the entire data pattern) can differ from the readouts of the control reference. Although a random explanation is theoretically possible, e.g., the control readouts were not representative and only rare “unlucky” outcomes, the confirmation by independent experiments points to a non-random cause. However, the reasons for this phenomenon are usually unclear, with biological effects and technical issues discussed (Krebs et al., 2018). If ignored, a control normalization can not only suggest false treatment-related effects at non-effective concentration ranges, but also contradict the meaning of a comparable relative effect scale and therefore invalidate a BMR: if non-effective concentrations produced normalized effect responses which are more than 10% different from the negative controls, then a regression modelling cannot establish a BMC10. To overcome the problem of misleading control data it has been suggested to estimate the control reference directly from the responses of the test concentrations, assuming that they can provide sufficient evidence for “non-exposure related endpoint activity”, which is often translated as equal assay responses at the lowest test concentrations over a sufficient large concentration range (Krebs *et al*., 2018). A refined control reference can then either be estimated directly from these data responses or from response estimated by concentration-response regression analysis, and then be used to re-normalize the entire data set such that the refined control reference is set to 100%.

The experimental design for the assays from the DNT IVB was chosen such that the lowest test concentrations were expected to produce no treatment-related effect responses. For more than 90% of all data sets the three lowest concentrations provided non-distinguishable effect responses, which we deemed as sufficient for using the control re-normalization on all data, and it was implemented in the automatic data evaluation pipeline as part of the standard protocol. This was mainly motivated by the frequent occurrence of misleading negative control responses observed for some of the assay endpoints.

On this background it is not very surprising that the choice of the normalization method not only led to very different BMC estimations but often completely failed, as documented by the number of BMRnr’s (Figure 5B): the intended BMR was not covered by control-normalized responses, and as consequence the regression analysis suggested a best fit model that also did not cover the BMR and therefore could not estimate a BMC. However, a re-normalization of the effect scale guaranteed coverage of the BMR, and accordingly the regression modelling was able to establish a BMC. Figure 7B provides another example for a gross data misinterpretation: normalized effect responses suggest a BMC for inhibition, re-normalized effect data a BMC for induction. Although the majority of data sets did not necessarily require a control-renormalization, a change to the standard control normalization still changed the hit category for approx. 5% of all endpoint-specific DNT hazard classifications (Figure 8D), and impacted all performance parameters about the DNT IVB’s predictivity negatively (Figure 9).

A re-normalization should only be applied if sufficient data evidence is provided to do it, otherwise an existing exposure effect can be judged wrongly as technical or biological artifact and misused as zero effect response in the statistical concentration-response analysis. This decision making is only difficult to resolve in an automatized HTS data evaluation and we cannot fully rule out that a re-normalization was wrongly used for some data sets. Therefore, we recommend for the future a list of criteria such that those data scenarios can be identified and flagged for an expert decision. Potential criteria for assuring a successful data re-normalization could be a certain minimum magnitude between the original and re-defined control references based on statistical reasoning, a minimum number of test low concentrations for which the effect responses provide no indications for a positive or negative trend and a minimum concentration range at which no effect can be judged. In case a control normalization can neither be judged by an automatic ruling system nor by an expert we suggest as last solution a repetition of the experiment at lower test concentrations.

### 4.3 Concentration-response regression model

A BMC is derived from a mathematical function which is fitted as parametric regression model to the experimental concentration-response data. As no unique mechanistic model exists that would allow a 100% accurate representation of every possible shape of a concentration-response pattern, a regression model is considered as empirical and judged as suitable only if it describes the data in the best possible way. Each model is characterized by a limited number of model parameters which provide the flexibility to fit the model as close as possible to the data: the more model parameters are considered the better the model will describe the data, and the more complex the observed data pattern is (e.g. non-monotony) the more model parameters are required to describe the data. However, the more model parameters are considered the more data evidence are required to support a reliable regression fit. If a model with too many parameters is chosen that is not supported by a sufficient amount of data (over-parametrization), the estimation method will result into overinflated model estimates and therefore in an unreliable BMC estimate with an extremely large CI (high difference between BLL and BUL). In the concentration-response plot, the latter is usually indicated by a huge “uneven” confidence belt around the regression fit with extreme peaks at concentrations at with the highest nonlinearity of the concentration-response pattern occurred. Moreover, two different models might describe the concentration-response data similarly well with identical BMC estimates, but the model with the lower number of model parameters will result into smaller confidence interval around the BMC (and which might therefore influence the hazard classification). Generally, the model with the lowest number of parameters is favored over a more complex model as long as it can describe the data almost as accurate as the more complex model (parsimony). The accuracy vs. parsimony trade-off is utilized by the AIC criterion in the model selection process of the best-fit method.

Data and the experimental design decide on which model can be chosen for a BMC estimation, and the minimal data requirements can be assigned directly to the model parameters: each parameter requires a specific data support from effect responses of at least one concentration, i.e. a model with five parameters would require five concentrations with effect responses that are specifically addressing the nature of the parameter. For instance, the model parameter describing the upper control asymptote (d in Table S3) requires that at least one concentration has produced responses that can be used as average control reference, the parameter describing the lower maximal asymptote (d in Table S3) requires that at least one concentration has produced responses that can be used as average control reference, and all other model parameters require data support from concentrations that neither have produced maximal or minimal responses. If these data requirements are not given, a BMC estimation should be considered always as unreliable from a strict statistical point of view. For example, the data example C in Figure 7 provides no effect responses between the minimal and maximal asymptote, and therefore a priori no regression model can be fitted to the data on sound statistical criteria. Nevertheless, the LL4 model (blue line) provides an BMC estimation including an 95% confidence belt which, in this case, appears to be well in line with the experimental data, and can be considered as sufficiently accurate for regulatory purposes as all three independent experiments produced nearly identical data outcomes.

A concentration-response function commonly used in pharmacology and toxicology is the Hill function, which is a reparametrized form of the 3-parameter log-logistic function (LL3, but without the log10 transformation of the concentration, Table S2). We chose this function as pre-defined model in comparison to the best-fit model approach since it a popular approach for regression of strict monotonic concentration-response patterns in comparable software (*tcpl*, Filer *et al*. 2016), and since it is also recommended by the OECD for continuous data (EFSA Scientific Committee, 2017). In line with theoretical expectations and previously reported simulation studies (Zhu *et al*., 2007; West *et al*., 2012; Piegorsch *et al*., 2013), the best-fit model approach responded more flexible to data sets and therefore resulted often to BMC estimations that differed significantly from those derived by the Hill model (Figure 5C). As a consequence, the sole application of the Hill model occasionally prevented the estimation of a BMC and its uncertainty, and therefore led to less data sets for which a hazard identification could be performed. This strongly suggests the use of several regression models in a best-fit approach, including functions with maximal two model parameters for “data-poor” sets and more complex functions for “data rich” scenarios.

Model averaging is historically motivated by the typically small number of doses in animal studies that can provide meaningful data for the regression modeling, and the subsequent problem that different regression models can describe the observed dose-response data equally well but interpolation in a dose region with little or no data may result into very different response (and BMD) estimates (EFSA, 2017). A statistical argument in favor of model averaging is that uncertainty of the model selection process of the best fitting method is not incorporated in the BMD and associated BMDL estimation (West et al., 2012). Our study shows no big differences between both methods, and we attribute the higher number of failed BMC estimates for model averaging (Figure 5D) due to the fact that the models with the lowest number of model parameters were not included in the pool of candidate models for model averaging: the 2-parameter exponential model (Table S2) was selected as best-fit model in approx. 33% of all model decisions (standard protocol), indicating that the data sets did not allow the selection of a more complex model with 3 model parameters, and as consequence model averaging did fail. It demonstrates that simple regression models are essential for “poor data” scenarios, i.e., data sets where maximally two concentrations responded with significant but often weak assay responses.

### 4.4 Uncertainty estimation of the BMC

Not only the BMC estimation is crucial for the hazard classification but also a correct derivation of its uncertainty, usually expressed as lower and upper confidence interval (CI). We have used various statistical methods which are implemented in the *drc* and *bmd* R package (delta approximation, inverse regression, resampling methods), and investigated how they can impact the hazard classification. It should be noted that these methods do not change the BMC estimation but try to calculate the uncertainty of the BMC estimation from the estimated regression model and experimental data. All methods have their pros and cons with different requirements to the data and regression models, and none of them can a priori be ruled out as inappropriate for the BMC estimation of a DNT IVB data set. Assuming that the correct regression model was chosen and the estimation method led to only one reliable BMC estimation, we used the inverse regression as CI reference for a best-fit model estimated BMC, and compared it to the ones derived from the delta method and parametric bootstrapping. The advantage of the inverse regression method is that the confidence of a regression curve can easily be assessed by a non-statistician by showing the concentration-response data together with the regression fit and its associated confidence interval in a common plot.

As shown in Figure 6H, different CI methods often resulted in largely different outcomes, with a moderate impact on the hazard classification (Figure 8D) and corresponding DNT IVB performance parameters (Figure 9). The delta method provides a means to estimate the approximate variance and CI of a model function when the function consists of one or more estimated model parameters, and where there is an estimate for the variance of each model parameter, with both derived from the successful fit estimation. The method implemented in the *drc* package (Ritz *et al*., 2015) is based on a first order approximation for the variance of the BMC, and thus expected to be accurate only for concentration-response pattern that show a minor non-linearity (Zhu, Wang and Jelsovsky, 2007; Moerbeek, Piersma and Slob, 2004). Higher order approximations of the delta method would be progressively more flexible and provide a better description of the BMC uncertainty but are currently not implemented. Therefore, it is not surprising that the delta approximation often failed with an unreliable CI spanning the entire range of test concentrations (Figure 7D), especially for the 2-parameter exponential function with a concentration term that is not log10-transformed. Based on the study outcome we deem this method as unfit for an automatic HTS data evaluation.

Whereat the delta method is entirely based on the outcomes from the regression fit and therefore provides a quick and easy way to calculate the IC for a BMC, resampling methods use only the regression model(s) and BMC estimation and develop the BMC uncertainty entirely from re-doing the regression analysis and BMC derivations on a huge number of concentration response data resampled from the original experimental data sets. This method puts strong emphasis on a “representative” data set for the resampling, and if violated, it is prone to biased interval estimations (i.e. mode of the resampled BMC distribution differs from the original BMC estimation) or, in worst-case, the simulations lead to an interval that hardly mirrors the observed data variability. Typically for DNT IVB, endpoints often produced responses with a relatively high between-study variability (documented in the corresponding BMRs, Table 1), with only a small sample size for resampling (3-5 experimental replicate medians), and with often only two or less test concentrations which provided significant responses distinguishable from the controls. These data scenarios are not optimal for regression resampling, and therefore it is not surprising that bootstrapping often resulted in very different, too wide confidence belts compared to those from inverse regression, or even completely failed (Figure 7E). To some extent this might also explain the different outcomes for model averaging, which was performed always in combination with bootstrapping.

Until generally applicable decision rules about the minimal data requirements for bootstrapping can be implemented in an automatic data evaluation platform, it is only difficult for the non-expert to make decisions about the usefulness of resampling for a particular data scenario. Therefore, our advice is that the user should have some experience regarding statistical resampling, or, if applicable, use inverse regression or related methods.

### 4.5 The choice of the BMR on the Hazard identification

The optimal choice of an endpoint-specific BMR level is always a comprise between “as close as possible” to the control reference (i.e. a BMC estimation as low as possible) and the statistical demands for providing a reliable BMC estimation for as many data sets as possible. In principle, each data set has its own optimal BMR, mainly defined by its between-experimental data variability and how well it can support the regression analysis. A BMR which guarantees a BMC for all future data sets would need to be chosen from the data set with the largest observed between-experimental data variability, and by this a relatively large BMR would be favored, with response levels around 50% for some of the assay endpoints (i.e. a BMC would equal an EC50 or IC50). However, a larger BMR leads to a higher BMC, and the consequence for all data sets which a much lower data variability is that their substance responses observed at concentration ranges below the BMC are ignored, and therefore contradict the intended regulatory meaning of a benchmark concentration. But more important, it would also rule out those data sets for a BMC estimation where the observed maximal responses are below the BMR (and thus a BMC cannot be established). The latter was decisive for ruling out many sets for a BMC estimation after we increased the BMRs by 20% in our standard protocol (e.g., changing BMC30 to BMC50, Figure 5E), and as consequence our simulations resulted into 5% different hazard classifications, with a change mainly from “hit” to “no hit” (Figure 8D). Therefore, the use of the most common descriptor for concentration response data in pharmacology and in vitro toxicology, an IC50 or EC50, cannot be recommend as surrogate for a BMC for endpoints of the DNT IVB.

A critical aspect is how a BMR can be derived: we used the 1.5 sigma rule, with sigma estimated as standard deviation from the between-experimental variation from a large set of historical data sets (Masjosthusmann et al., 2020). For a sample size of 3-5 independent experiments, we expected for the majority of data sets the estimation of a BMC if a true BMC was present in the data, but nevertheless our standard protocol might have failed to identify a hazard because the BMR was selected as too low for a particular data set. Our study outcomes do not provide the exact number how often this might have happened as it would require for each individual data set a statistical power analysis and corresponding estimation of the detection limit, however, assuming that the scatter between the experimental replicate medians always followed the Gaussian distribution we expect this to be the case in less than 1% of all cases.

### 4.6 Hazard identification and software

A huge number of free software packages for the statistical analysis of dose-response data and dose-response modelling are available, with PROAST (RIVM National Institute for Public Health and the Environment), BMDS (US EPA), ToxCast pipeline (tcpl, Filer et al. 2017) or BMCeasy (Krebs et al., 2019) just to mention a few. Similar to the R packages we use (drc and bmd, Ritz et al., 2015 and Jensen et al., 2020), most of these software packages provide a variety of options in order to respond as flexible as possible to the various data scenarios a user can possibly face, and as consequence, always a minimum of statistical knowledge is demanded from the user. Similar to the tcpl pipeline we became interested in an automated data evaluation platform with no required user intervention and addressing the specific features of DNT data or other data from organotypic cultures. To our experience, the proposed standard protocol is for an automated data evaluation pipeline the best compromise between the various statistical methods without “overcomplicating” the regression analysis and the corresponding BMC estimation. The drawback of an automated analysis is always the danger of not being prepared to deal with an unusual data set, a scenario that most likely can only be avoided by analysing each data set individually by an expert. The strength of our data evaluation platform is the integration of endpoint-specific hazard classifications, including flagging systems for uncertain cases, which none of the software packages mentioned above offer. We consider it crucial for the hazard assessment to differentiate between general cell toxicity and specific DNT hits.

### 4.7 Conclusion

The comparative study between various statistical methods involved in the estimation of a BMC and its associated uncertainty for a huge number of concentration response data sets from the DNT IVB revealed the following main conclusions:

1. The normalization of effect data to the outcomes of test concentrations can be a viable option to safeguard against an ill-defined negative control reference and therefore avoid a biased BMC estimation and incorrect hazard alerts. This re-normalization of response data should be done whenever sufficient data evidence is provided for non-exposure related effect responses at lowest concentrations and which have been confirmed by independent experiments. Optimally it should be decided on a case-by-case basis by the experimenter, and more efforts are required to integrate decisions for a re-normalization in automatic data evaluation routines.
2. The pool of candidate models for the parametric regression analysis should include as simple as possible mathematical functions in order to enable a BMC estimation for data sets which provide only little data support for the regression modelling. This can be either the exponential or linear function, with both including maximally only two model parameters.
3. Simple common statistical methods such as the delta method do not necessarily guarantee a reliable estimation about the uncertainty of a BMC confidence and depend strongly on the chosen regression model and its non-linearity close to the BMR. In contrast, more sophisticated methods such as resampling require more data support which is often not given by the experiments. Invers regression provided the best way to judge a BMC uncertainty.
4. Data sets with only two or less effective concentrations are often borderline to a reliable statistical analysis, but nevertheless provide sufficient data for a “pragmatic” solution. The BMC for the specific DNT endpoints of these data sets is usually at high concentrations and within (or close) to the cytotoxic concentration ranges, and thus are most likely not too be classified as “specific hit”.
5. The BMR level should be chosen as close as possible to the control level without compromising the statistical concentration-response analysis. Setting it too high (e.g. 50%) involves the danger of overlooking hazard responses which can lead to erroneous hazard hit classifications.
6. An endpoint-driven hazard classification method is essential for a reliable identification of hazard alerts, and DNT-specific endpoints should always take general cell health into account. The automatized data evaluation should include a decision making that pinpoint to data scenarios which require a manual expert judgment for the hazard classification.

Although this study was conducted on concentration response data from only the DNT IVB, we think many of the conclusions can be generalized to data from other specific toxicological endpoints, especially in the rising field of organotypic/stem cell-based cultures. It demonstrates that statistical decisions which seem to be of minor importance can become decisive if it comes to the hazard classification of a test substance. It also demonstrates how important fit-for-purpose, internationally harmonized and accepted data evaluation and analysis procedures are for an objective hazard classification.

## Author contribution

All authors read, commented, and approved the manuscript. **Hagen Eike Keßel**: study conception, data analysis, software development, supervision, figure design, writing of article. **Stefan Masjosthusmann**: study conception, figure design, supervision. **Kristina Bartmann**: investigation. **Jonathan Blum**: investigation. **Arif Dönmez**: software development, data analysis. **Nils Förster**: software development, data analysis. **Jördis Klose**: investigation. **Axel Mosig**: software development, supervision. **Melanie Pahl**: investigation. **Marcel Leist**: supervision. **Martin Scholze**: supervision, software development, data analysis, writing of article. **Ellen Fritsche**: study conception, supervision, funding acquisition, project administration, writing of article.

## Acknowledgements

The authors are grateful to Katie Paul Friedmann (US EPA) for data integration and transfer to the ToxCast data base. We would like to thank Signe Marie Jensen (University of Copenhagen) for her help with proper application of the *drc* and *bmd* R packages for data analysis. This work was supported by the European Food Safety Authority (EFSA - Q - 2018 – 00308), the Danish Environmental Protection Agency (EPA) under the grant number MST-667-00205 and the project CERST (Center for Alternatives to Animal Testing) of the Ministry for culture and science of the state North-Rhine Westphalia, Germany (file number 233-1.08.03.03-121972/131 – 1.08.03.03 – 121972).

