## Supplementary Data for "Biostatistics and its impact on hazard characterization using in vitro developmental neurotoxicity assays"

### Impact of biostatistical data evaluation methods on hazard characterization using the neurosphere model as case study

#### Supplementary Data

### 1 Supplementary material and methods

#### 1.1 Pre-processing of endpoints

**Table S1: Pre-processing of endpoints**
Pre-processed endpoints (left) are calculated with raw endpoints (right).

| Pre-processed endpoint | Formular (raw endpoints) |
|---|---|
| neuronal differentiation [120h] | $\frac{\text{number of neurons}}{\text{number of all cells}}$ |
| oligodendrocyte differentiation [120h] | $\frac{\text{number of oligodendrocytes}}{\text{number of all cells}}$ |
| migration distance neurons [120h] | $\frac{\text{migration distance of neurons}}{\text{migration distance of all cells}}$ |
| migration distance oligodendrocytes [120h] | $\frac{\text{migration distance of oligodendrocytes}}{\text{migration distance of all cells}}$ |
| Viability UKN4 | $\frac{\text{number of selected objects}}{\text{number of valid objects}}$ |
| Viability UKN5 | $\frac{\text{number of selected objects}}{\text{number of valid objects}}$ |

#### 1.2 Regression models

Table S2 shows all parametric regression functions that defined the pool of candidate models for the best fit method. All models were applied by the *drm* function of the *drc* package with all key parameters (in the following written in cursive) set as follows: *formular* was given by a list of averaged replicate values and corresponding dose values, *type* was set to “continuous” for continuous data regression, *robust* was set to “mean” for least-square-estimation of continuous data, *fct* was set to one of the regression models listed in the drc syntax column of Table S2.

**Table S2: Regression models**

| Model[2)] | *drc* syntax | Model equation[1)] |
|---|---|---|
| general logistic | logistic2() | $f(x) = c + \frac{d - c}{(1 + \exp(b(\log(x) - \log(e))))^f}$ |
| 3-parameter log-logistic | LL.3() | $f(x) = 0 + \frac{d - 0}{1 + \exp\big(b(\log(x) - \log(e))\big)}$ |
| 4-parameter log-logistic | LL.4() | $f(x) = c + \frac{d - c}{1 + \exp\big(b(\log(x) - \log(e))\big)}$ |
| 2-parameter exponential | EXD.2() | $f(x) = 0 + (d - 0)(\exp\left(-\frac{x}{e}\right))$ |
| 3-parameter exponential | EXD.3() | $f(x) = c + (d - c)(\exp\left(-\frac{x}{e}\right))$ |
| 3-parameter Weibull | w1.3() | $\mathrm{f(x) = 0 + (d - 0)exp(-exp\big(b(log(x) - e)\big))}$ |
| 4-parameter Weibull | w1.4() | $\mathrm{f(x) = c + (d - 0)exp(-exp\big(b(log(x) - e)\big))}$ |

[1)] The parameters b, c, d, e of the model equations are estimated by concentration-response analysis: d is always estimated as model asymptote for the negative control response, c describing maximal responses at high concentrations (lower model asymptote), c and e provide flexibility in describing the location and steepness of the concentration-response pattern.

[2)] Model name and abbreviation from *Analysis of Dose-Response Curves* (Ritz et al. 2016).

#### 1.3 Classification model

**Table S3: Specific, unspecific and viability-related endpoint correlations for the classification model**
Specific endpoints are shown with their affiliated unspecific and viability-related endpoints. To gain DNT specific classifications, DNT specific endpoints are compared to either one or two unspecific endpoints (measuring general cell health). If an effect was detected in one of the affiliated viability-related endpoints, only cytotoxicity endpoints were used as reference.

| Specific Endpoint | Unspecific Endpoint 1 | Unspecific Endpoint 2 | Viability-related Endpoint 1 | Viability-related Endpoint 2 |
|---|---|---|---|---|
| migration distance radial glia [72h] | cytotoxicity (migration) [72h] | | | |
| migration distance radial glia [120h] | viability (migration+differentiation) [120h] | cytotoxicity (migration+differentiation) [120h] | migration distance radial glia [120h] | cell number [120h] |
| cell number [120h] | viability (migration+differentiation) [120h] | cytotoxicity (migration+differentiation) [120h] | migration distance radial glia [120h] | cell number [120h] |
| proliferation by BrdU [72h] | viability (proliferation) [72h] | cytotoxicity (proliferation) [72h] | | |
| proliferation by area [72h] | viability (proliferation) [72h] | cytotoxicity (proliferation) [72h] | | |
| neuronal differentiation [120h] | viability (migration+differentiation) [120h] | cytotoxicity (migration+differentiation) [120h] | migration distance radial glia [120h] | cell number [120h] |
| oligodendrocyte differentiation [120h] | viability (migration+differentiation) [120h] | cytotoxicity (migration+differentiation) [120h] | migration distance radial glia [120h] | cell number [120h] |
| migration distance neurons [120h] | viability (migration+differentiation) [120h] | cytotoxicity (migration+differentiation) [120h] | migration distance radial glia [120h] | cell number [120h] |
| migration distance oligodendrocytes [120h] | viability (migration+differentiation) [120h] | cytotoxicity (migration+differentiation) [120h] | migration distance radial glia [120h] | cell number [120h] |
| neurite length [120h] | viability (migration+differentiation) [120h] | cytotoxicity (migration+differentiation) [120h] | migration distance radial glia [120h] | cell number [120h] |
| neurite area [120h] | viability (migration+differentiation) [120h] | cytotoxicity (migration+differentiation) [120h] | migration distance radial glia [120h] | cell number [120h] |
| Migration UKN2 [24h] | Viability UKN2 [24h] | | | |
| Neurite Area UKN4 [24h] | Viability UKN4 [24h] | | | |
| Neurite Area UKN5 [24h] | Viability UKN5 [24h] | | | |

**Table S4: Alerts used to flag classification data for manual evaluation**
During endpoint classification, the data is checked for uncertainties and an alert is produced, if the resulting classification has high uncertainty. The according alerts (left) with reasoning (right) are shown.

| Alert | Explanation |
|---|---|
| BMCU above concentration testrange | If the upper confidence limit of the BMC is above the tested concentration range, it has a high uncertainty and automated classification cannot be made. |
| high CI width | A high CI width indicates uncertainty in the BMC estimation and should therefore be checked for the final classification. A CI width was considered as high, if BMCU/BMCL > (BMC*5)/(BMC/5). |
| no data for significance | Statistical significance could not be calculated (likely due to sample size of n<=2), but is needed for classification. |
| Issues with predict CI | Algorithmic failure did not allow the calculations of the BMC upper or lower limit. Therefore, no automated classification can be made. |
| no data | Necessary data for the classification was missing. |

### 2 Supplementary results

#### 2.1 Assay performance of all control compounds

To assess how changes in the data evaluation protocol might impact the evaluation of the DNT IVB's predictivity, 28 reference chemicals of known DNT and 17 negative control chemicals were selected (Masjosthusmann et al. 2020), with all 45 substances tested in the DNT IVB. A negative compound was considered as true negative (abbreviated as "TN" in the table below) if it was not classified as specific hit or borderline in any of the assays. Else, it was considered as false positive (FP). A positive compound was considered as true positive (TP), if it was classified as specific hit or borderline in at least one assay. Else, it was considered as false negative (FN).

**Table S5: Assay performance for negative controls**
Negative controls (n=17) used to determine the specificity of different protocols. Negative control compounds are listed in the first column. Remaining columns represent the compound classifications for each protocol used in this study. Specificity is given as true negative rate at bottom row and is calculated as the percentage of true negatives (TN) from all negative controls.

| Negative controls | Standard Protocl | Replicate Mean | Control-Normalized | LL3rm | Boot-strapping | Model Averaging | BMR30+50 |
|---|---|---|---|---|---|---|---|
| **Amoxicillin** | TN | TN | TN | TN | TN | TN | TN |
| **Aspirin** | TN | TN | FP | TN | TN | FP | TN |
| **Buspirone** | TN | TN | TN | TN | TN | TN | TN |
| **Chlorpheniramine maleate** | TN | FP | TN | TN | TN | FP | TN |
| **D-Glucitol** | TN | TN | TN | TN | TN | TN | TN |
| **D-Mannitol** | TN | TN | TN | TN | TN | TN | TN |
| **Diethylene glycol** | TN | TN | TN | TN | TN | TN | TN |
| **Doxylamine succinate** | TN | TN | TN | TN | TN | TN | TN |
| **Famotidine** | TN | TN | TN | TN | TN | TN | TN |
| **Ibuprofen** | TN | TN | TN | TN | TN | TN | TN |
| **Metformin** | TN | TN | TN | TN | TN | TN | TN |
| **Metoprolol** | TN | TN | TN | TN | TN | TN | TN |
| **Penicillin VK** | TN | TN | TN | TN | TN | TN | TN |
| **Saccharin** | TN | TN | TN | TN | TN | TN | TN |
| **Sodium benzoate** | TN | FP | TN | TN | TN | TN | TN |
| **Warfarin** | TN | TN | TN | TN | TN | TN | TN |
| **Specificity (True Negative Rate in %)** | **100.0** | **87.5** | **93.8** | **100.0** | **100.0** | **87.5** | **100.0** |

**Table S6: Assay performance for human positive controls**

Human positive controls (n=9) used to determine the sensitivity of different protocols. Human positive control compounds are listed in the first column. Remaining columns represent the compound classifications for each protocol used in this study. Sensitivity is given as true positive rate at bottom row and is calculated as the percentage of true positives (TP) from all human positive controls.

| Human positives | Standard Protocol | Replicate Mean | Control-Normalized | LL3rm | Boot-strapping | Model Averaging | BMR30+50 |
|---|---|---|---|---|---|---|---|
| 2,2',4,4'-Tetrabromodiphenyl ether | TP | TP | TP | TP | TP | TP | TP |
| Cadmium chloride | TP | TP | TP | TP | TP | TP | TP |
| Chlorpyrifos | TP | TP | FN | TP | TP | TP | TP |
| Dexamethasone | TP | TP | TP | TP | TP | TP | TP |
| Hexachlorophene | TP | TP | TP | TP | TP | TP | TP |
| Lead(II) acetate trihydrate | TP | TP | TP | TP | TP | TP | TP |
| Manganese(II) chloride | TP | TP | FN | TP | TP | TP | FN |
| Methylmercury(II) chloride | TP | TP | TP | TP | TP | TP | TP |
| PBDE 99 | TP | TP | TP | TP | FN | FN | TP |
| Sensitivity (True Positive Rate in %) | 100.0 | 100.0 | 77.8 | 100.0 | 88.9 | 88.9 | 88.9 |

**Table S7: Assay performance for *in vivo* positive controls**

*In vivo* positive controls (n=19) used to determine the sensitivity of different protocols. *In vivo* positive control compounds are listed in the first column. Remaining columns represent the compound classifications for each protocol used in this study. Sensitivity is given as true positive rate at bottom row and is calculated as the percentage of true positives (TP) from all *In vivo* positive controls.

| *in vivo* positives | Standard Protocol | Replicate Mean | Control-Normalized | LL3rm | Boot-strapping | Model Averaging | BMR30+50 |
|---|---|---|---|---|---|---|---|
| (+-)-Ketamine hydrochloride | FN | FN | FN | FN | FN | FN | FN |
| (-)-Nicotine | FN | FN | FN | FN | FN | FN | FN |
| 5,5-Diphenylhydantoin | FN | FN | FN | FN | FN | FN | FN |
| Acrylamide | TP | TP | TP | TP | TP | TP | TP |
| Chlorpromazine hydrochloride | TP | TP | TP | TP | TP | TP | TP |
| Deltamethrin | TP | TP | TP | TP | TP | TP | TP |
| Domoic acid | FN | FN | FN | FN | FN | FN | FN |
| Haloperidol | TP | TP | TP | TP | TP | TP | TP |
| Heptadecafluorooctanesulfonic acid potassium salt | TP | TP | TP | TP | TP | TP | TP |
| Maneb | TP | FN | TP | TP | TP | FN | FN |
| Methylazoxymethanol acetate | TP | TP | TP | TP | TP | TP | TP |
| Paraquat dichloride hydrate | TP | TP | TP | TP | TP | TP | TP |
| Perfluorooctanoic acid | FN | FN | TP | FN | FN | FN | FN |
| Sodium valproate | TP | TP | TP | TP | TP | TP | TP |
| Tebuconazole | TP | TP | TP | FN | TP | TP | TP |
| Tributyltin chloride | TP | TP | TP | TP | TP | TP | TP |
| Trichlorfon | TP | TP | TP | FN | TP | TP | TP |
| Triethyltin bromide | TP | TP | FN | FN | TP | FN | TP |
| all-trans-Retinoic acid | TP | TP | TP | TP | TP | TP | TP |
| Sensitivity (True Positive Rate in %) | 73.7 | 68.4 | 73.7 | 57.9 | 73.7 | 63.2 | 68.4 |
